# Assessing the chemistry and bioavailability of dissolved organic matter from glaciers and rock glaciers

**DOI:** 10.1101/115808

**Authors:** Timothy Fegel, Claudia M. Boot, Corey D. Broeckling, Ed K. Hall

**Affiliations:** Natural Resource Ecology Laboratory, Colorado State University; U.S. Forest Service, Rocky Mountain Research Station; Department of Chemistry, Colorado State University; Proteomics and Metabolomics Facility, Colorado State University, Department of Ecosystem Science and Sustainability, Colorado State University

## Abstract

As glaciers thaw in response to warming, they release dissolved organic matter (DOM) to alpine lakes and streams. The United States contains an abundance of both alpine glaciers and rock glaciers. Differences in DOM composition and bioavailability between glacier types, like rock and ice glaciers, remain undefined. To assess differences in glacier and rock glacier DOM we evaluated bioavailability and molecular composition of DOM from four alpine catchments each with a glacier and a rock glacier at their headwaters. We assessed bioavailability of DOM by incubating each DOM source with a common microbial community and evaluated chemical characteristics of DOM before and after incubation using untargeted gas chromatography mass spectrometry based metabolomics (GC-MS). Prior to incubations, ice glacier and rock glacier DOM had similar C:N ratios and chemical diversity, but differences in DOM composition. Incubations with a common microbial community showed DOM from ice glacier meltwaters contained a higher proportion of bioavailable DOM (BDOM) and resulted in greater bacterial growth efficiency (BGE). After incubation, DOM composition from each source was statistically indistinguishable. This study provides an example of how MS based metabolomics can be used to assess effects of DOM composition on differences in bioavailability of DOM. Furthermore, it illustrates the importance of microbial metabolism in structuring composition of DOM. Even though rock glaciers had significantly less BDOM than ice glaciers, both glacial types still have potential to be important sources of BDOM to alpine headwaters over the coming decades.

**Key Points:** Bioavailability of organic matter released from glaciers is greater than that of rock glaciers in the Rocky Mountains.

The use of GC-MS for ecosystem metabolomics represents a novel approach for examining complex organic matter pools.

Both glaciers and rock glaciers supply highly bioavailable sources of organic matter to alpine headwaters in Colorado.

## 1. Introduction

Ice glaciers (hereafter glaciers) are perennial ice structures that move and persist in areas where annual snow accumulation is greater than annual snow ablation at decadal or longer time spans. Rock glaciers are flowing bodies of permafrost, composed of coarse talus and granular regolith both bound and lubricated by interstitial ice (Berthling, 2011). Both glaciers and rock glaciers integrate atmospherically deposited chemicals with weathering products and release reactive solutes to adjacent surface waters (Williams et al., 2007; Dubnick et al., 2010; Fellman et al., 2010; Stibal et al., 2010; Stubbins et al., 2012). On average, alpine glaciers appear to be responsible for the release of a larger flux of carbon (0.58 +/− 0.07 Tg C) from melting ice annually when compared to continental glaciers (Hood et al., 2015). Dissolved organic matter (DOM) from large alpine glaciers of the European Alps and Alaska has been shown to be bioavailable and thus can support microbial heterotrophy (Hood et al., 2009; Singer et al., 2012). In the contiguous United States, both glaciers and rock glaciers, each with distinct geophysical attributes (Fegel et al. 2016), are common to many alpine headwaters, with rock glaciers being an order of magnitude more abundant than glaciers. However, little is known about the quantity or quality of DOM being released from these smaller glaciers and rock glaciers in mountain headwater ecosystems of the western U.S. (Woo et al., 2012; Fegel et al., 2016).

Assessment of the bioavailability of aquatic DOM to date has largely relied on bioassays that measure the rate and amount of DOM consumed over time (e.g. Amon and Benner, 1996; del Giorgio and Cole, 1998; Guillemette and del Giorgio, 2011). Bioassays present difficulties in assessing DOM consumption because each assessment requires an independent incubation, creating the potential for confounding effects due to differences in the microbial communities among incubations. Broad molecular characterizations of DOM (Benner, 2002; Berggren and del Giorgio, 2015) have also previously been applied to estimate bioavailable DOM (BDOM), though these analyses often only address differences in bioavailability of either coarse categories of organic molecules or individual model compounds rather than the composition of the DOM pool as a whole (e.g., del Giorgio and Cole, 1998). Recently the research community has begun to explore high resolution analytical chemistry to assess the molecular characteristics of environmental DOM (e.g., Kujawinski, 2002; Kellerman et al., 2014, 2015; Andrilli et al., 2015). Whereas no single method can identify the entire spectrum of compounds present in an environmental DOM pool (Derenne and Tu, 2014), mass spectrometry (MS) has the ability to define structural characteristics of thousands of individual compounds contained within a single DOM sample.

There is a suite of challenges in trying to concurrently understand the composition and bioavailability of DOM. Difficulties in experimentally linking the molecular characterization of DOM pools to the proportion of BDOM are partly due to the highly diverse constituent compounds that compose natural DOM and the inherent challenges in characterizing that diversity (Derenne and Tu, 2014). In addition, different microbial communities may access different components of the same DOM pool making it challenging to compare how composition affects bioavailability among ecosystems. However, even in the face of these challenges some broad patterns of the relationship between the composition and bioavailability of DOM are beginning to emerge. For example, studies from glacial, estuarine, and marine environments have shown a clear correlation between proteinaceous DOM and a high proportion of BDOM (Andrilli et al., 2015). Understanding which constituent molecules influence the bioavailability of DOM is an important step in understanding how DOM pools contribute to heterotrophy among aquatic ecosystems and how different components of DOM alter the residence time of that DOM pool in the environment.

We hypothesized that differences in the origin of DOM between glaciers and rock glaciers would result in differences in molecular composition of DOM between glacier types with the potential to alter the proportion of BDOM. DOM derived from glaciers originates from *in situ* microbial activity via phototrophy on the glacier surface, and byproducts of heterotrophic bacterial activity in the glacier porewaters and subsurface (Anesio et al., 2009; Hood et al., 2009; Singer et al., 2012; Fellman et al., 2015; Fegel et al., 2016). In addition, atmospheric deposition of organic matter onto the glacier surface can be an important source of DOM to the glacier meltwaters (Stubbins et al., 2012). DOM from rock glaciers is an amalgam of compounds originating both from multicellular vegetation growing on the rock glacier surface and microbial processing within the rock glacier itself (Wahrhaftig and Cox, 1959; Williams et al., 2007). Previous research has shown that DOM released from glaciers and rock glaciers in the western United States differ in their optical properties, with glaciers having higher amounts of protein-like DOM than rock glaciers, as identified through fluorescence spectroscopy (Fegel et al., 2016).

In this study, we assessed differences in the composition of DOM between glaciers and rock glaciers and asked which of those differences (if any) correspond to differences in the proportion of BDOM from each glacier type. We also asked whether the composition and bioavailability of DOM in glacier and rock glacier meltwaters in the western United States is similar to what has been reported for other glacial meltwaters globally. Here we present the results of incubations of DOM with a common microbial community from four pairs of glacier and rock glacier meltwaters (eight DOM sources in total). We used GC-MS with a nontargeted metabolomics workflow to define differences in DOM from glaciers and rock glaciers before and after each incubation. In addition to measuring bacterial respiration during each incubation, we measured change in bacterial cell numbers, allowing us to also assess how DOM from each glacial source influenced bacterial production relative to bacterial respiration.

## 2. Methods

### 2.1 Site Description

Paired glaciers and rock glaciers from four watersheds on the Front Range of Northern Colorado were selected based on their size (>0.5km^2^) and the proximity to each other within the watershed (Figure 1). We collected samples of glacier meltwater in September 2014 to maximize the contribution from ice melt and minimize annual snowmelt contribution (Figure 1, Table 1). A complete description for each site is provided in Fegel et al. (2016).

**Figure 1.**
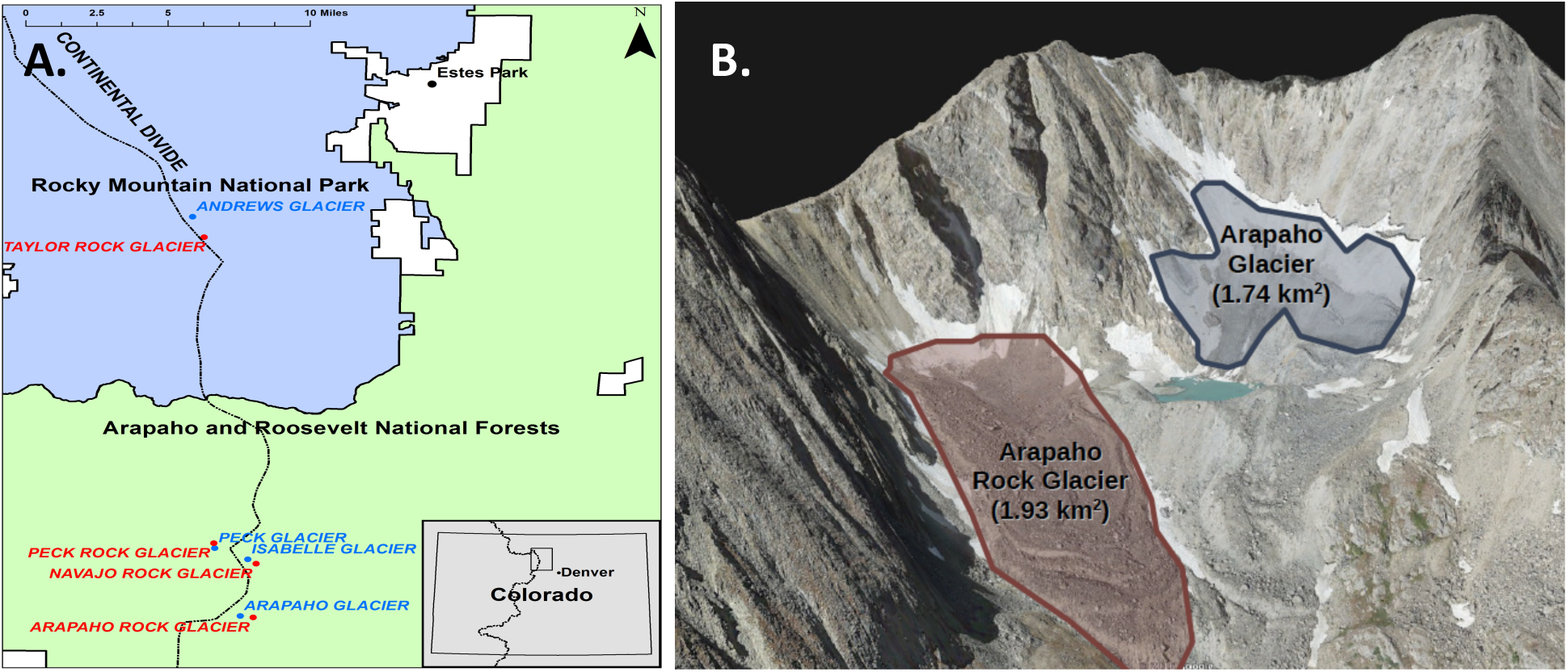
Site Map A) Locations of each glacier type that was sampled for this study within the state of Colorado. In total four pairs of glaciers and rock glaciers were sampled along the Front Range of Colorado. **B) Arapaho Glacier and Arapaho Rock Glacier** form a pair of a glacier and a rock glacier from the same watershed and are shown here to illustrate the differences between the two types of glaciers.

**Table 1.**
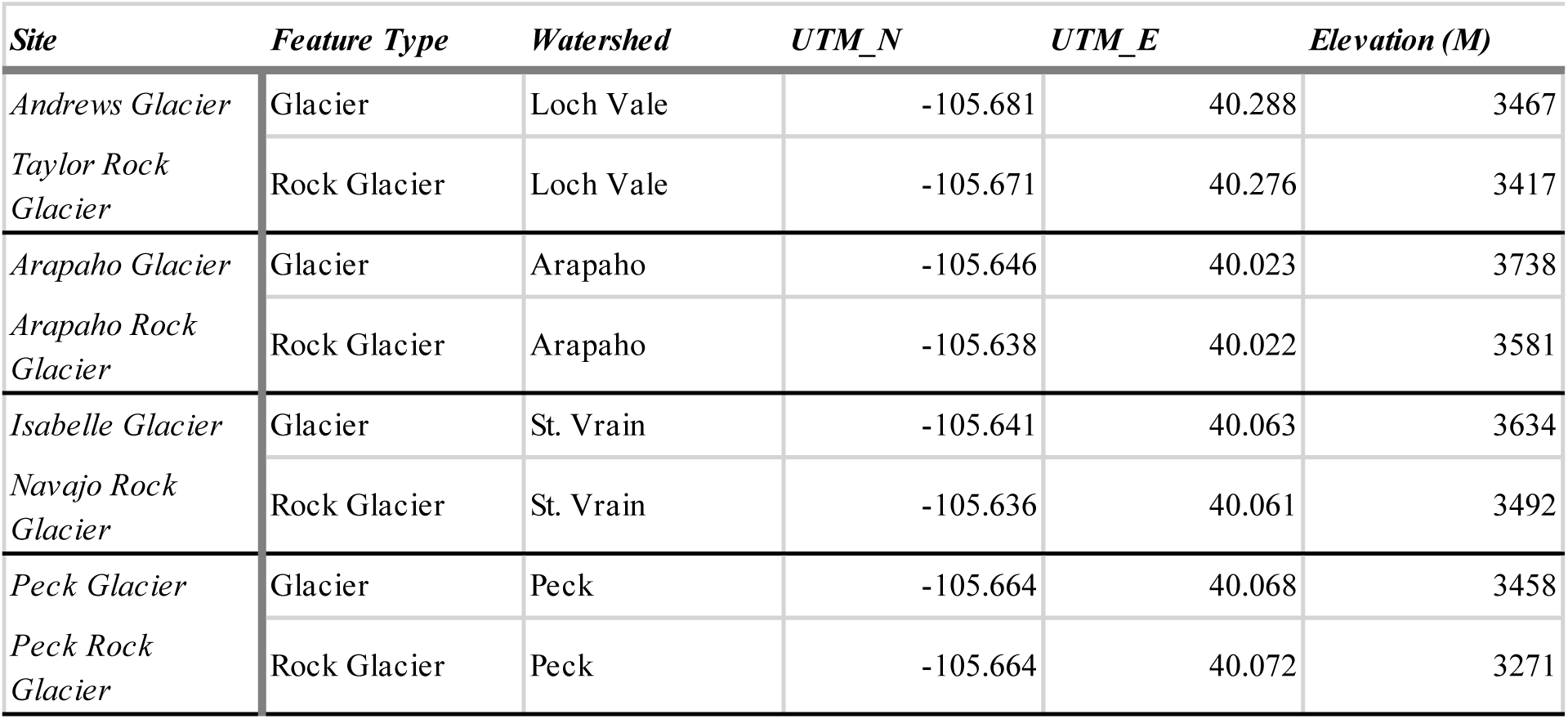
Site description for the four pairs of glaciers and rock glaciers sampled within our study. Italicized names represent non-official names of glaciers and rock glaciers in study. Coordinates are given in Universal Transverse Mercator North (UTM_N) and Universal Transverse Mercator East (UTM_E).

### 2.2 Field extraction of DOM

Meltwaters from each glacier type were collected in the early to midmorning (05:00-10:00) to minimize diurnal variability in ice melt among glaciers. DOM was concentrated in the field from 20 L of meltwater collected at the terminus of each glacier type using a slightly modified protocol from Dittmar et al. (2008). Briefly, meltwater samples were passed through precombusted (450° C, 5hr) Whatman GF/F filters (GE Whatman, Pittsburg, PA, USA), acidified to ~ pH 2 with 32% HCl, and concentrated using Bond Elut PPL carbon extraction cartridges (Bond Elut-PPL, 500 mg 6 mL Agilent, Santa Clara, CA, USA).

Concentrated DOM (0.67-1.75 mg C, Supporting Information Table S3) on each PPL cartridge was eluted in the laboratory with HPLC grade MeOH and collected into cleaned, combusted, and preweighed 120 mL borosilicate bottles. DOM was isolated from MeOH in each bottle by running clean N_2_ gas over the open samples until all MeOH was evaporated. Samples were kept at room temperature. A portion of the dried DOM was redissolved in water and used for the reactivity assays, the remaining DOM was re-eluted into MeOH for metabolomic analysis via GC-MS.

### 2.3 DOM bioavailability experiments

Dried DOM samples from each of the eight sites were diluted to 4 mg L^−1^ C with MilliQ (>18MΩ H2O) and incubated in vitro with a natural bacterial community collected from The Loch (−105.6455, 40.2976), a small subalpine lake in Rocky Mountain National Park, CO, USA. Unfiltered lake water collected from The Loch was aged for ~2 years at 5 °C to remove the majority of the BDOM. At the start of the experiment DOM concentration of the aged lake water was 0.7 mg C L^−1^, with a pH of around 6.7. The goal of the bioassay was to determine which portions of the glacial DOM pools could be consumed by a naturally occurring bacterial community. Aging the lake water allowed us to reduce the DOM contribution from the lake water and at the same time provide a common community of naturally occurring aquatic alpine microbes that should not be preferentially disposed to glacier or rock glacier DOM as The Loch is located in a watershed containing both glacier types and at a similar distance from both. Prior to incubation, 2L of aged lake water was filtered through a precombusted (450° C, 5hr) Whatman GF/C filter (1.2μm nominal pore size) to remove bacterial grazers (e.g. protists and metazoans). At the initiation of the experiment (t=0), three aliquots (10 mL) of filtered, aged lake water were preserved with 2% formalin (final concentration, from 37% formaldehyde) and set aside for enumeration of bacteria. A second set of three aged lake water aliquots (7 mL) was analyzed for initial DOM (as dissolved organic carbon) and total dissolved nitrogen (TDN) on a Shimadzu TOC-VWS analyzer (Shimadzu Corp., Kyoto, Japan).

Normalized concentrations of DOM (4 mg L^−1^ C) were created in each incubation bottle by adding 60.96 mL of filtered aged lake water, between 3.80 and 9.04 mL of concentrated glacial DOM solution (depending on the initial carbon concentration) and filling the remaining volume to 70 mL total volume with MilliQ water (>18MΩ H2O). Total MilliQ water additions added to each microcosm ranged between 0 and 5.2 mL. This resulted in concentrations of ~4 mg L^−1^ C in each incubation bottle for all DOM sources (Table S1). Experimental controls contained only GF/C filtered lake water with no added DOM. An analytical control of MilliQ water was also included to correct for any instrumental drift that occurred during the experiment.

All microcosms were incubated simultaneously at 15 °C. To calculate microbial respiration, we measured changes in dissolved oxygen (DO) at 1 minute intervals in each microcosm using an Oxy-4 fiber optic dissolved oxygen probe for at least six hours, every other day (PreSens, Precision Sensing GmbH, Regensburg, Germany). The incubation was terminated at 10 weeks, when the fastest metabolizing microcosm approached 4 mg L^−1^ DO to avoid the potential for anaerobic metabolism to contribute to carbon consumption. All measurements that were below the recommended detection value of the Oxy-4 mini were removed because of the potential for inaccurate estimations of DO (PreSens Oxy-4 mini Instruction Manual, Precision Sensing GmbH, 2011). Absolute values from the raw fiber optic measurements were corrected for analytical drift of the probe by subtraction of changes in signal from the MilliQ water analytical control over the course of the experiment.

### 2.4 Bacterial cell counts and bulk chemistry

From each microcosm we collected a 2 mL aliquot preincubation (described above) and post incubation and preserved each with 2% formalin (final concentration) to assess changes in cell abundance during the course of the incubation. Aliquots were filtered onto 0.2 μm Millipore polycarbonate filters and stained with Acridine Orange for enumeration, following the methods of Hobbie et al. (1977; Supporting Information Data Set S1).

### 2.5 Metabolite analyses

Prior to incubation each sample of DOM was prepared for metabolomic analysis by redissolving a portion of the dried DOM into fresh HPLC grade methanol to a final concentration of 2 mg C mL^−1^. Post incubation DOM samples were collected separately from each individual microcosm (n=16). Each 58mL (70mL minus the 2mL bacterial cell count aliquot and the 10mL post incubation DOC/TDN subsample) sample was filtered through preleached 0.45 *μ*m Millipore filters. The sample was freeze dried, and the remaining DOM was weighed and redissolved into HPLC grade methanol to a final concentration of 2 mg mL^−1^ C for GC-MS analysis. Carbon concentration of DOM assays was measured before and after incubation (Table S2 and Table S3). We tested the preleached, blank Millipore filters for carbon and found a negligible contribution (Table S2). Dissolved carbon values are accurate within +/-5% using a nonpurgeable dissolved organic carbon method (EPA method 415.1, Shimadzu Corporation, Columbia, MD). Percent recovery of DOM was calculated by comparing the carbon content of the normalized freeze dried sample to the actual values of the unconcentrated meltwater for each glacier type and averaged 71% +/− 17% (Table S3). To increase the volatility of molecules analyzed through GC-MS, samples for metabolomics analysis were derivatized with trimethylsilane (TMS) using standard protocols (Text S1: Derivatization Process for GC-MS, Pierce, 1968).

### 2.6 Metabolomics

Both preincubation and post incubation DOM samples were analyzed with inline gas chromatography mass spectrometry (GC-MS) at the Proteomics and Metabolomics Facility at Colorado State University. Metabolites were detected using a Trace GC Ultra coupled to a Thermo ISQ mass spectrometer (Thermo Scientific, Waltham, MA, USA). Samples were injected in a 1:10 split ratio twice in discrete randomized blocks. Separation occurred using a 30 m TG-5MS column (0.25 mm i.d., 0.25 μm film thickness, Thermo Scientific, Waltham, MA, USA) with a 1.2 mL min^−1^ helium gas flow rate, and the program consisted of 80°C for 30 sec, a ramp of 15°C per min to 330°C, and an 8 min hold. Masses between 50-650 m/z were scanned at 5 scans sec^−1^ after electron impact ionization (Broeckling et al., 2014).

### 2.7 Data analysis

#### 2.7.1 Metabolite analysis

For each sample, a matrix of molecular features defined by retention time and ion mass to charge ratio (*m/z*), was generated using feature detection and alignment in the XCMS R package. We used an in house clustering tool, RAMClustR, to group ions belonging to the same compound based on coelution and covariance across the full dataset (Broeckling et al., 2014). Spectral abundance (hereafter, abundance) equates to the weighted mean for the peak area of each ion that clustered to a single compound. Compound annotation was prioritized based on order of spectral abundance and differentiation between glacier types. Annotated compounds are those with spectral similarity to reference spectra in an MS library (Sumner et al. 2007). Annotation confidence levels 1-4 are used to indicated the certainty of the structural assignment. Level 1 annotations are reserved for identified compounds which are compared with data from identical acquisition parameters for authentic reference standards, and level 4 annotations are unknown compounds. Unknown compounds retain compound names generated by the clustering step and have the form C#, e.g. C40. We used in house, NISTv12, Golm, Metlin, and Massbank metabolite libraries for annotation of TMS derivatized molecules. Annotated compounds had high similarity (>90%) to spectra from known standard compounds within the databases used (Figure S1).

Compound molecular rank was calculated by ordering DOM compounds by their average abundance preincubation versus postincubation. Chemical diversity was calculated using the Shannon-Wiener diversity index (Shannon, 1948) by treating each unique DOM compound as a ‘species’, similar to method used by Kellerman et al. 2014 with the Chao1 index for chemical diversity. Shannon-Wiener was calculated for DOM composition both before and after incubation to estimate changes in chemical diversity through microbial metabolism. Chemical superclasses for the most abundant annotated compounds, as well annotated compounds that were significantly different by glacier type or incubation period were assigned using conversion of chemical name to InChI Code using the Chemical Translation Service (http://cts.fiehnlab.ucdavis.edu), and then the InChI Code was converted to the chemical taxonomy based on molecular structure using ClassyFire (http://classyfire.wishartlab.com, Feunang et al., 2016). Compound superclasses are the second level of hierarchical classification in the chemical taxonomy and consist of 26 organic and 5 inorganic groups. Compounds with the same superclass share generic structural features that describe their overall composition or shape. For example, the organic carbon superclass contain at least one carbon atom excluding isocyanide/cyanide and their non-hydrocarbyl derivatives, thiophosgene, carbon diselenide, carbon monosulfide, carbon disulfide, carbon subsulfide, carbon monoxide, carbon trioxide, carbon suboxide, and dicarbon monoxide; whereas the organic oxygen superclass contains at least one oxygen atom; the organic nitrogen superclass contains at least one nitrogen atom, and the benzenoid superclass has aromatic compounds containing one or more benzene rings. A full description for each superclass in the dataset has been included in the supplemental information.

#### 2.7.2 Analysis of DOM bioavailabilitys

To estimate the size of the bioavailable DOM pool (BDOM) and the recalcitrant DOM pool we fit oxygen consumption rates to a Berner-Multi-G two-pool decay model using SAS (Berner, 1980; Guillemette and del Giorgio, 2011). Oxygen consumption was averaged for each glacier type and confidence intervals were calculated at *α*=0.05. Dissolved oxygen curves generated from the incubation were fit to the equation: 

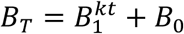

Where *B*_T_ is the total carbon pool converted from measured moles of consumed oxygen to moles of carbon, with the assumption of a respiratory quotient (RQ) of 1. B_1_ is the BDOM carbon pool, k is the decay rate constant of the BDOM pool, t is time, and B_0_ is the recalcitrant carbon pool. We used a Wilcoxon signed rank test to compare statistical differences between glacier and rock glacier B_1_ and B_0_ pool sizes. Total carbon consumed was calculated as the difference in preincubation and post incubation measured DOM carbon concentrations. Many recent studies have compared different BDOM models and found that multi-g models can overestimate BDOM pool size over the long term and that DOM bioavailability is a continuum better represented by beta and gamma distribution shapes (Vahatalo et al. 2010; Vachon et al. 2016; Mostovaya et al. 2017). However, these studies were based on incubations lasting longer than a year and in study systems having relatively small BDOM pools. Further, these reactivity continua were used to estimate total pool sizes at watershed scales. For this study we chose the multi-g model because of the relatively short incubation period (10 weeks), and the hypothesized large BDOM pool size of glaciers. Use of the multi-g model allowed us to examine the relative sizes of BDOM versus more recalcitrant DOM pools throughout the course of our incubation length.

In addition to evaluating the BDOM and DOM pools from each glacier we assessed how the microbial community metabolized each sample by calculating bacterial growth efficiency (BGE) of the bacterial community. Bacterial production was measured as the change in cell number over time and converted to units carbon using an estimate of 20 fg C per bacterial cell (Borsheim and Bratbak, 1987; Data Set S1). To calculate BGE we divided bacterial production by the sum of bacterial respiration and bacterial production (del Giorgio and Cole, 1998).

#### 2.7.3 Statistical analyses

Differences in DOM composition between paired treatments (glaciers vs. rock glaciers), and pre/post incubation were compared with ANOVA. Mixed model ANOVA was performed using the lme4, lmerTest, and lsmeans packages in R. The model was specified: site + type + incubation, with watershed and injection replicate set as random variables. The mixed model was applied to relative intensities of individual GC-MS identified compounds and p-values for all statistical tests were adjusted for false positives using the Bonferroni-Hochberg method in the p.adjust function (R Core Team, 2014). Common MS contaminants were removed from the dataset prior to further processing (Keller et al., 2008). We visualized differences in DOM composition by glacier type pre and post incubation using PCA conducted on mean centered and pareto scaled data using the pcaMethods package (R Core Team, 2014). PCA analyses were computed between glacier types preincubation, and a separate PCA was computed for the pre/post incubation comparison. Identification of the compounds correlated with separation in ordination space was conducted in two steps, first ANOVAs on each of the PC scores (with the anova(aov) function), for each sample was used to determine which axes best describe the experimental treatments (Figure 2). This approach provided information on how well the PCs describe the variability of the dataset with respect to the experimental design, without compromising the advantages of an unsupervised analysis. The experimental treatments may not account for the majority of the variability in the dataset (i.e. described by ordination along PC1 and PC2), such that PCs best describe the experimental conditions may have PC higher numbers (e.g. PC5 for glacier type, Figure 2). The second step to determine which compounds contributed disproportionately to separation by glacier type was an outlier test (pnorm function and p value fdr adjustment for multiple testing) performed on the loadings values for each PC (Figure 2). In addition to these analyses we also calculated average fold changes (the ratio of the pre incubation to post incubation intensity) of specific compounds indicated as important in the univariate and multivariate analyses. We used the nonparametric Wilcoxon signed rank test for each glacier pair to compare differences in BGE, C:N, chemical diversity, and change in chemical diversity between glacier types as these variables did not meet the expectations of equal variance and normal distribution required by parametric tests.

**Figure 2.**
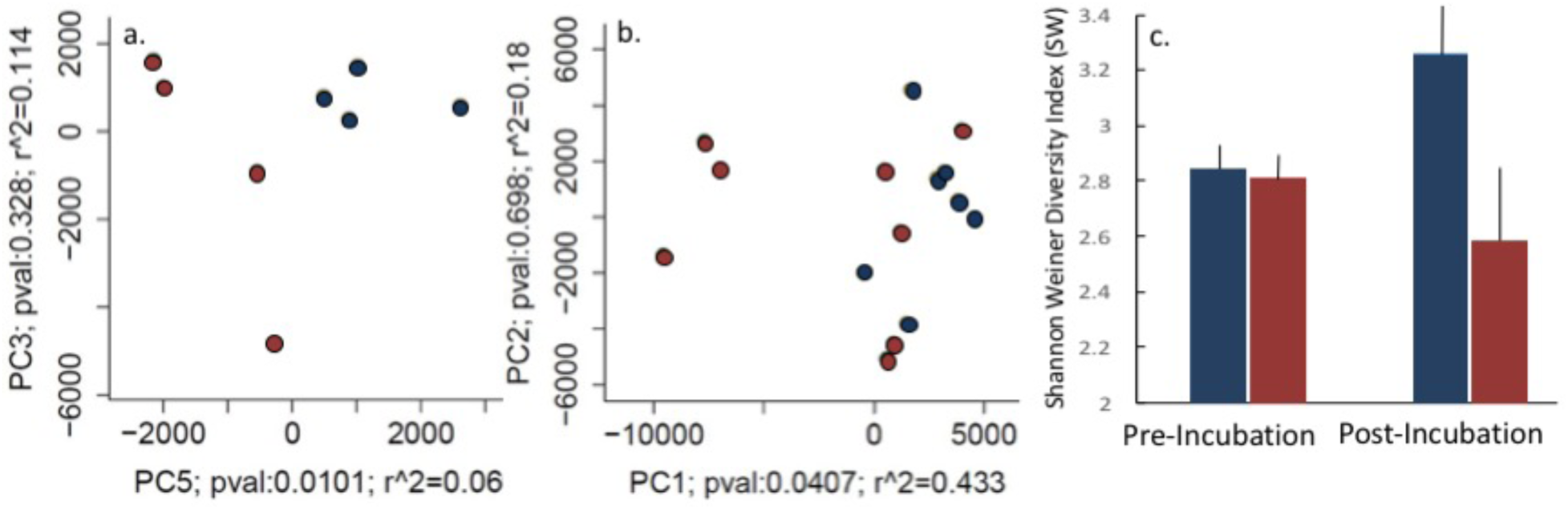
PCA analysis by glacier type for DOM compounds identified using GC-MS. **a)** PCA plots showing separation along PC5 between glaciers (blue) and rock glaciers (red) before incubation and **b)** no significant difference between glacier types after incubation (p-value of 0.0.407 and 0.0681 for PC1 and PC2, respectively). **c)** The Shannon-Wiener Index (SW) for chemical diversity was similar between glaciers and rock glaciers before incubation, however microbial metabolism increased chemical diversity in glacier DOM and decreased chemical diversity in rock glacier DOM. Statistical analysis of differences between glacier types can be found in Table 3.

**Table 2.** Characteristics of DOM from glaciers (G) and rock glaciers (RG). ***DOC_initial_*** is in initial concentration of carbon (mg C L^−1^) from the unconcentrated meltwater collected from each glacier. C:N is the ratio of carbon to nitrogen for preincubation meltwater DOM. ***Δ***D.O (change in dissolved oxygen) and ***Δ***DOM (change in dissolved organic carbon concentration as mg C L^−1^) represent the change in concentration of each during the course of the incubation (mg O_2_ L^−1^), SW = Shannon Wiener Diversity Index, and ΔSW = change in SW of DOM before and after the incubation. Bold values represent significant differences between glaciers and rock glaciers (p<0.01) using the Wilcoxon signed rank test for paired nonparametric samples. Standard deviations of mean values are listed in parentheses.

|  | Isabelle<br>Glacier | Peck<br>Glacier | Andrews<br>Glacier | Arapaho<br>Glacier | Peck<br>Rock<br>Glacier | Navajo<br>Rock<br>Glacier | Arapaho<br>Rock<br>Glacier | Taylor<br>Rock<br>Glacier | Mean<br>Glacier | Mean Rock<br>Glacier |
| --- | --- | --- | --- | --- | --- | --- | --- | --- | --- | --- |
| $DOC_{initial}$<br>($mg\ C\ L^{-1}$) | 0.78 | 0.93 | 0.82 | 1.08 | 0.98 | 1.21 | 0.42 | 1.1 | 0.90 (0.13) | 0.93 (0.35) |
| $C:N$ | 1.65 | 2.02 | 3 | 2.73 | 2.63 | 1.48 | 2.45 | 0.86 | 2.35 (0.62) | 1.85 (0.83) |
| $\Delta DO$<br>($mg\ O_2\ L^{-1}$) | 6.03 | 3.95 | 4.11 | 3.97 | 3.52 | 2.81 | 4.12 | 2.69 | **4.51 (1.02)** | 3.29 (0.67) |
| $\Delta DOM$<br>($mg\ C\ L^{-1}$) | 2.5 | 1.89 | 2.64 | 2.84 | 2.09 | 1.71 | 3 | 2.25 | **2.47 (0.41)** | 2.26 (0.54) |
| $SW$ | 2.92 | 2.89 | 2.68 | 2.84 | 2.8 | 2.83 | 2.79 | 2.8 | 2.83 (0.11) | 2.80 (0.02) |
| $\Delta SW$ | 0.19 | 0.88 | 0.13 | 0.52 | 0.54 | -0.32 | -1.1 | -0.01 | **0.43 (0.30)** | 0.22 (0.59) |

**Table 3.**
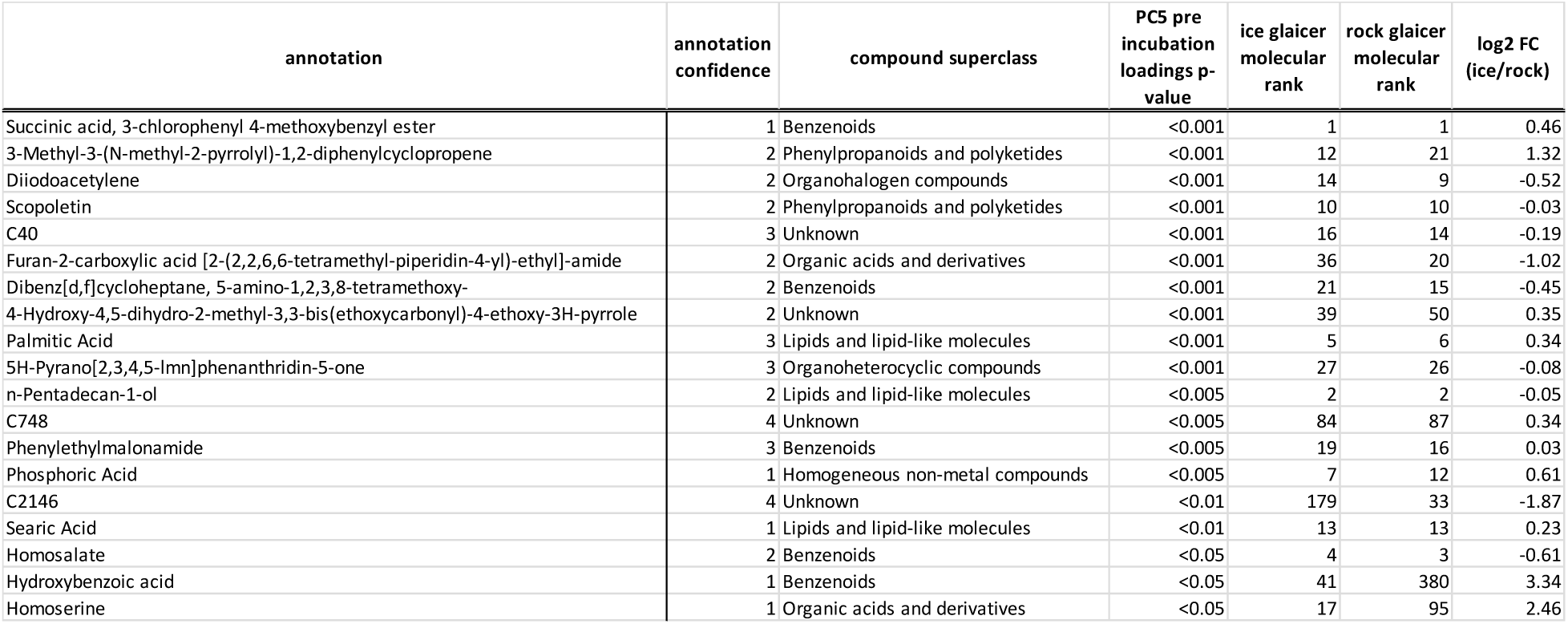
**DOM constituents that drove the differences between glaciers and rock glaciers before incubation**, as identified with multivariate analysis (ANOVA on PC axis 5 loadings). Annotation confidence is noted with 2D structure assignment using a reference standard from an in-house library as a score ‘1’ through a structurally unknown feature of interest of a confidence level ‘4’ (Blazenovic 2018). Positive values for Log2 fold change (FC) indicate greater relative abundance in glaciers versus rock glaciers, and negative Log2 FC values indicate greater relative abundance in rock glaciers versus glaciers (Supporting Information Data Set S2).

## 3. Results

DOM had C:N ratios around 2 (2.35+/-0.62 for glaciers, 1.85+/-0.83 for rock glaciers) for both glacier types, which is consistent with the elevated inorganic nitrogen reported from glaciers in Colorado and elsewhere (Williams et al., 2006; Barnes et al., 2014; Fegel et al., 2016, Saros et al., 2010; Slemmons et al., 2013). Prior to incubation neither the concentration, chemical diversity or the C:N ratio of the DOM were significantly different between glaciers and rock glaciers (Table 2).

### 3.1 Glacier and rock glacier DOM metabolomics

The composition of glacier and rock glacier DOM consisted of 2,056 compounds after ordinary contaminates from mass spectrometry analysis were removed (Data Set S2). Of those 2,056 compounds each individual compound consisted of 3 to 170 individual mass spectral features, for a total of 14,571 mass spectral features in the dataset. The univariate analysis (paired mixed model ANOVA) did not identify any compounds that had significantly different abundances between glacier types (ice vs. rock) before incubation. However, the mixed-model univariate analysis did indicate 889 compounds that were significantly different before and after incubation with the common microbial community (Figures 3, 6, and Data Set S2). The mixed-model also indicated an additional 21 compounds had a significant interaction term for the glacier type by incubation period interaction (Data Set S2). Of the total 2,056 compounds, we chose 322 for annotation based either on high abundance (n=303) or disproportionate contribution to separation between glacier types in multivariate space (n=19, Table 3).

Eighty-three percent of the variation in pre-incubation DOM composition (i.e., differences in abundance of 2056 compounds) could be expressed by the first two principal components. However, glacier type did not separate along these axes (ordination not shown). Rather differences in DOM composition between glacier type was clearly visible in the space defined by the 5^th^ principal components (Figure 2A). An additional multivariate analysis (outlier test on PC loadings) indicated 19 compounds that contributed disproportionately to multivariate separation along PC axis 5 which explained 18% of the variance in the data set and best described separation between glacier and rock glacier DOM (Figure 2A). Of the 19 compounds that drove separation by glacier types (Table 3) we made 16 annotations, with 3 compounds unable to be annotated in the current mass spectral reference libraries. Prior to incubation, glaciers and rock glaciers had similar representation among the 8 compound superclasses in the data set (Figure 3).

**Figure 3:**
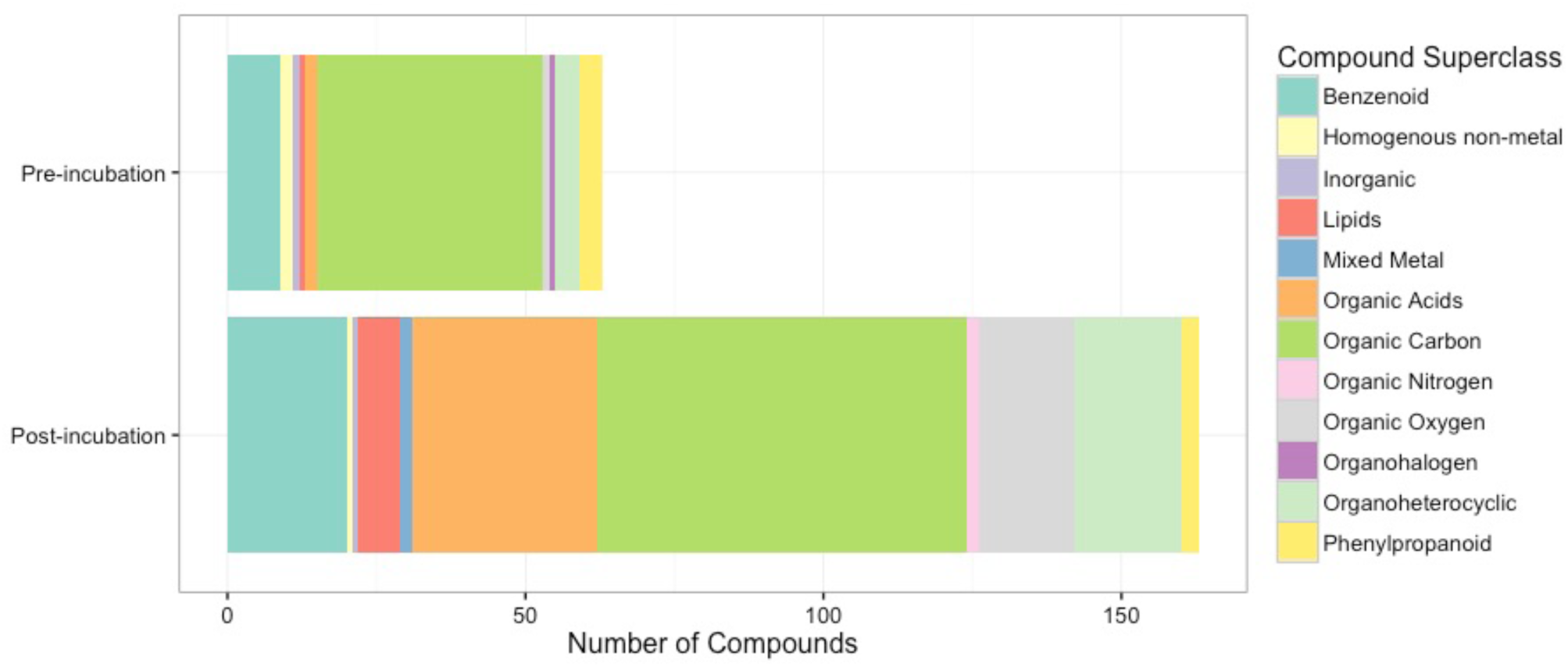
Annotated compounds that were significantly different pre and post incubation grouped by compound superclasses. Microbial metabolism during the incubation resulted in an increased total number of annotatable compounds with large increases in the representatives in organic acid, organic carbon, organic oxygen, and heterocyclic compound superclasses during the incubation.

We next evaluated the rank abundance of individual compounds within each DOM pool. The top 25 most abundant molecules were similar between glacier types prior to incubation and were primarily from chemical superclasses including benzenoids, phenylpropanoids, and polyketides (Data Set S2).

### 3.2 Incubations

Dissolved oxygen measurements from all incubations were fit to a two pool decay model that allowed us to estimate the initial rate of respiration (*k*), the size of the BDOM pool (B_1_), and recalcitrant DOM pool (B_2_), from each DOM source. The initial bacterial respiration rates (*k*) were not significantly different between glacier types (Figure 4, Table 4). However, a significantly larger portion of glacier DOM was bioavailable (B_1_ = 66.0 ± 10.6%) compared to rock glacier DOM (B_1_= 46.4 ± 8.9%, p <0.01, Table 4). BGE was also greater for bacterial communities incubated with glacier DOM compared to rock glacier DOM (BGE_G_ = 0.26 ± 0.13, BGE_RG_ = 0.16 ± 0.16, Table 4).

**Figure 4:**
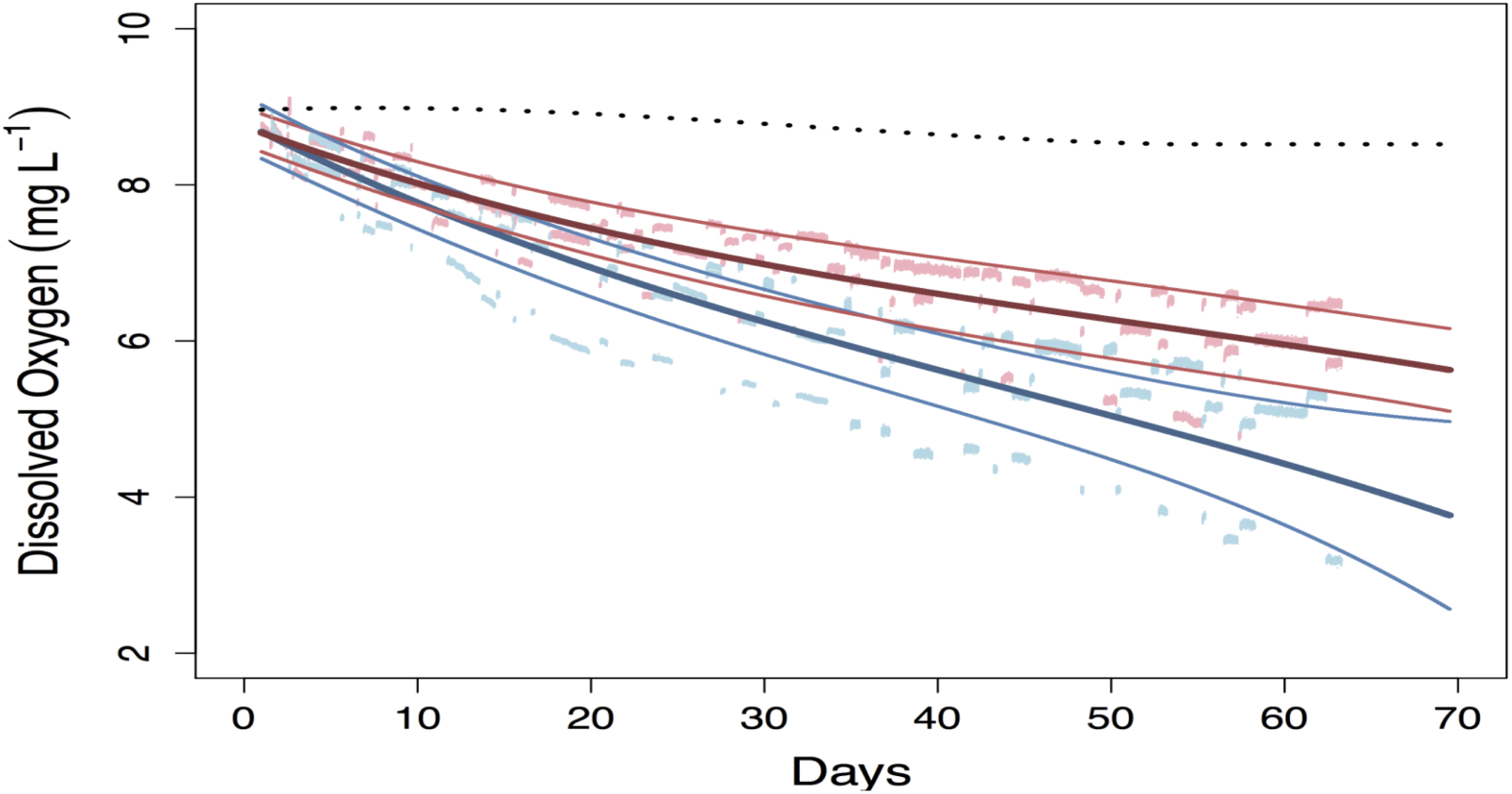
**Results from our laboratory incubation of DOM** from four different glaciated watersheds on the Front Range of Colorado. Here, values are averaged for each of the four glaciers (blue) and rock glaciers (red) and smoothed using a third order polynomial regression function (R^2^=0.999). 95% Confidence intervals are shown in light blue solid lines for glaciers and in pink solid lines for rock glaciers. The dotted black line is the experimental control (aged lake water with microbial community but no additional DOM).

**Table 4.** Results from carbon decay model where B_O_ represents the size of the recalcitrant pool of DOM (mg C L^−1^), B_1_ is the size of the BDOM pool (mg C L^−1^), *k* is the decay constant (mg C L^−1^ h^−1^), and BGE = bacterial growth efficiency. Glaciers had a larger percentage of BDOM compared to rock glaciers and rock glaciers had a larger proportion of recalcitrant carbon compared to glaciers. BGE was also higher for glaciers than rock glaciers. Bold values represent significant differences between glaciers and rock glaciers (p<0.01). Standard deviations of mean values are listed in parentheses.

|  | *Andrews*<br>*Glacier* | *Arapaho*<br>*Glacier* | *Isabelle*<br>*Glacier* | *Peck*<br>*Glacier* | *Arapaho*<br>*Rock*<br>*Glacier* | *Navajo*<br>*Rock*<br>*Glacier* | *Peck*<br>*Rock*<br>*Glacier* | *Taylor*<br>*Rock*<br>*Glacier* | *Mean Glacier* | *Mean Rock*<br>*Glacier* |
| --- | --- | --- | --- | --- | --- | --- | --- | --- | --- | --- |
| $B_0$ | 1.6 | 1.86 | 0.67 | 1.89 | 2.66 | 2.82 | 1.66 | 2.35 | **1.51 (0.54)** | 2.37 (0.63) |
| $B_1$ | 2.8 | 2.6 | 3.51 | 2.62 | 1.9 | 1.71 | 2.62 | 1.91 | **2.88 (0.53)** | 2.04 (0.56) |
| $k$ | 0.0003 | 0.0006 | 0.0008 | 0.0008 | 0.0019 | 0.0010 | 0.0007 | 0.0011 | 0.0008 (0.00) | 0.0010 (0.00) |
| % BDOM | 63.64 | 58.3 | 83.97 | 58.09 | 41.67 | 37.75 | 61.21 | 44.84 | **66.00 (10.61)** | 46.37 (8.93) |
| % Recalc. | 36.36 | 41.7 | 16.03 | 41.91 | 58.33 | 62.25 | 38.79 | 55.16 | 34.00 (7.03) | **53.63 (6.84)** |
| BGE | 0.37 | 0.39 | 0.39 | 0.3 | 0.13 | 0.06 | 0.13 | 0.07 | **0.263 (0.13)** | 0.157 (0.16) |

In the experimental control DOM decreased from 0.7 mg L^−1^ C at the beginning of the experiment to 0.16 +/− 0.08 mg L^−1^ C at the end. This change in DOM was ~6x less than the 3.06 +/− 0.23 mg L^−1^ C on average that was consumed during the course of the incubations for the microcosms amended with glacier or rock glacier DOM and should not have impacted the difference in BDOM observed between glaciers and rock glaciers (Tables S2 and S3). While carbon content did decrease slightly in the experimental control through the incubation, consumption of DOM in the experimental control was minor compared to both types of glacier incubations, and is mostly likely due to the change between the temperature the water was aged at (5 °C) and the temperature the experiment was conducted at (15 °C). This small change in DOM in the lake water controls is consistent with the small change in cell counts we observed of the lake water incubations (2.98*10^8^ change in cells) relative to cell changes in the microcosms that received glacier or rock glacier DOM (1.93*10^9^ average change in cells). Along with the minimal change in dissolved oxygen consumption (<0.5mg O_2_ L^−1^, dotted black line in Figure 4) these results suggest microbial activity was low in the lake water controls without amendments but was stimulated by the addition of glacier and rock glacier DOM (Data Set S1).

### 3.3 Changes in DOM during the incubation

The biggest differences in DOM composition were between pre- and post-incubation DOM pools. Incubations with the common microbial community altered the composition of the DOM from each glacier type, resulting in fewer compounds with high abundance and more compounds with low abundance (Figure 5). Univariate analysis indicated that 889 compounds differed significantly between the pre and post incubation DOM pools. Of those 889 compounds 419 had a greater abundance pre-incubation, whereas 469 had a greater abundance post incubation. We were unable to annotate the majority (~%75) of the 889 compounds that changed significantly during incubation (pre incubation 85% were unknown, and post incubation 65% were unknown). However, during the course of the incubation the number of compounds we were able to annotate increased 159% from 63 compounds pre-incubation to 163 compounds post-incubation. Of the compounds that could be annotated we observed a greater number of compounds from the organic acid, organic carbon, organic oxygen, and organoheterocyclic superclasses in the post incubation DOM pool compared to the pre-incubation DOM pool (Figure 3, Data Set S2).

**Figure 5.**
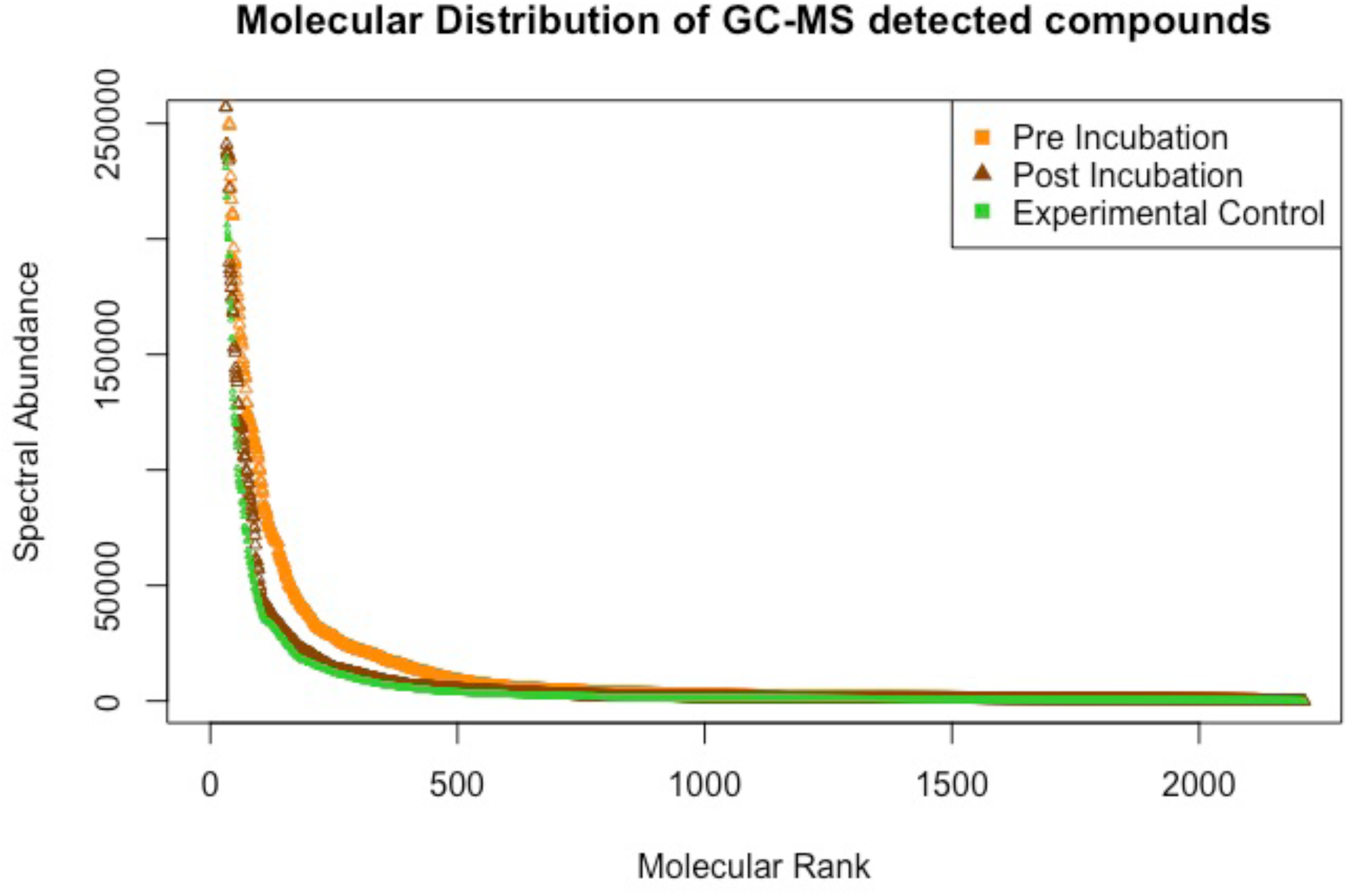
Molecular distribution of DOM compounds. Distribution of compounds by preincubation (orange) and post incubation (maroon) and experimental control (green, i.e., aged lake water). Many of the compounds present before incubation that were of intermediate abundance were metabolized during the incubation. The most abundant molecules were different between preincubation and post incubation metabolomics analysis (see Figure 6, Data Set S2).

Changes in the molecular composition of DOM during incubation restructured and reorganized the order of compounds within the molecular rank abundance curve of the post incubation DOM pools compared to the pre-incubation DOM pools (Figure 5, Data Set S2). This resulted in a reversal of compound molecular rank, with the compounds of highest relative abundance in the pre incubation samples having the lowest relative abundance in post-incubation DOM pools (Figure 6, Data Set S2). For example, homosalate ranked 3^rd^ in abundance pre-incubation and 563^rd^ post-incubation, and serine ranked 754^th^ pre-incubation and 5^th^ post-incubation. However, similar to how preincubation glacier and rock glacier DOM composition was analogous, post incubation glacier and rock glacier derived DOM shared the same most abundant compounds (Data Set S2).

**Figure 6:**
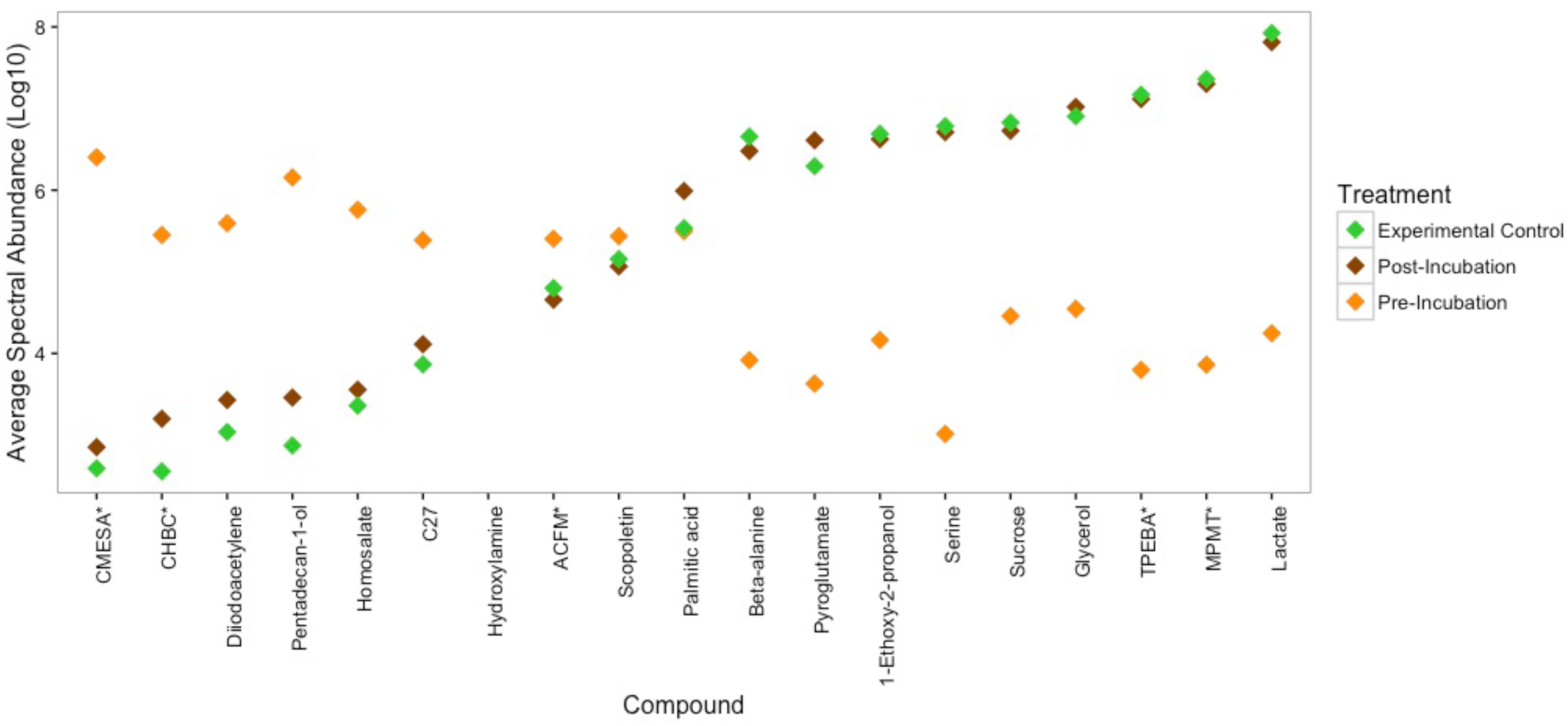
Change in the most abundant compounds pre versus post incubation. Microbial metabolism restructured the DOM pool, resulting in the most abundant compounds post incubation becoming more similar to the aged lake water control (*compound names abbreviated: ACFM = methanone, (2-amino=5chlorophenyl)(2-fluorophenyl), CHBC = p-Canophenyl 4’-heptyl-4-biphenylcarboxylate, CMESA = succinic acid, 3-chlorophenyl 4-methoxybenzyl ester, MPMT = 1,2,4-triazole, 3-mercapto-4-phenyl-5-methyl, TPEBA = benzenecarbothioic acid, 2,4,6-tritheyl-, S-(2-phenylethyl) ester).

Post incubation glacier and rock glacier DOM samples were more similar in structure and composition to that of the alpine lake DOM pool (i.e., the DOM of the experimental control) than compared to their pre-incubation DOM pools (Figures 5 and 6). This was true for both glacier and rock glacier DOM pools. Interestingly, chemical diversity of DOM between glacier types diverged during the course of the incubations. Post incubation glacial DOM had significantly higher diversity (SW = 3.26) compared to preincubation glacial DOM (SW=2.83), whereas post incubation rock glacier DOM had significantly lower diversity (SW=2.58) compared to preincubation rock glacier DOM (SW=2.80, Table 2, Figure 2C). This divergence in chemical diversity between DOM from each glacial type was due to an increase in the richness (number of compounds present) from preincubation to post incubation DOM pools (Figure 5, Data Set S2). While there were differences in the chemical diversity of glacier and rock glacier derived DOM post incubation, multivariate analysis (PCA) no longer indicated significant separation in the molecular composition of DOM between glacier types for any PC axes (Figures 2A and B).

## 4. Discussion

Our results demonstrate that chemically complex DOM released from both glaciers and rock glaciers supported bacterial respiration and bacterial secondary production. There were also significant differences in the size of the BDOM pools between glacier types, with glaciers having a higher proportion of BDOM than rock glaciers and supporting a higher BGE than rock glaciers. At the time of sampling, the composition of glacier and rock glacier DOM pools were not statistically distinct from one another, however 19 compounds correlated with separation in ordination space (Figure 2B). Processing DOM with a common microbial community resulted in a DOM pool with decreased chemical diversity of rock glacier DOM but increased chemical diversity of glacial DOM and more similar DOM composition between glacier types. After incubation both DOM pools had rank abundance of DOM compounds similar to that of the aged alpine lake water. These results suggest that microbial metabolism altered DOM to similar composition and structure, presumably reflective of the composite metabolism of the common microbial community.

### 4.1 Glaciers as a source of BDOM

The proportion of BDOM (66%) from glacier meltwaters in Colorado fell within the range of what has been reported in previous studies of glacial DOM in southeast Alaska (23-66%), the European Alps (58%), and the Tibetan plateau (46-69%, Hood et al., 2009; Singer et al., 2012; Spencer et al., 2014; Fellman et al., 2015). This suggests that on average glacier DOM around the world is enriched in BDOM relative to what has been reported for inland waters (4-34.5%) (Volk et al., 1997; Wiegner et al., 2006; Guillemette and del Giorgio, 2011). The mean amount of BDOM we report for rock glaciers (46%) was also higher than many inland waters, and within the range reported for other North American glaciers. Carbon concentrations in meltwaters from glaciers and rock glaciers in our study were low (0.5-1.5 mg C L^− 1^; Table 2), but again similar to previously reported mean glacial meltwater DOM concentration of 0.97 mg C L^−1^ (Dubnick et al., 2010; Stubbins et al., 2012; Singer et al., 2012; Hood et al., 2015). Knowing the volume of ice and estimates of discharge could allow us to estimate the total pool size of BDOM within glaciers and rock glaciers and an annual amount of BDOM released from each. Unfortunately, even with best available technologies, estimating discharge is exceedingly difficult due to the many shallow meltwater flow paths from both glacier types. Therefore, in this study we focused on differences between both the bioavailability and composition of DOM from each glacier type.

We suggest that differences in both the origin and delivery of DOM between glacier types in part explains why we saw higher proportions of BDOM in glaciers relative to rock glaciers. Rock glaciers host mosses, lichens, and vascular plants, including woody shrubs and trees (Wahrhaftig and Cox, 1959; Cannone and Gerdol, 2003; Burga et al., 2004). DOM released from these multicellular phototrophs include a wide variety of complex organic acids and polymers that are related to the secondary growth of the phototroph and are potentially difficult for microorganisms to metabolize (Wetzel, 1992; Rovira and Vallejo, 2002). Conversely, BDOM in glaciers is more likely to come from microbial production; either on the ice surface (Antony et al. 2017), in cryoconite holes (Sanyal et al., 2018), within the meltwater channels, or below the glacier (Fellman et al., 2009), and from atmospherically deposited materials (Stubbins et al., 2012). Organic molecules from microbial biomass or those small enough to be volatile should, on average, be more biologically available than plant derived organic matter, especially those compounds associated with the secondary metabolism of plants (Sun et al., 1997; Raymond and Bauer, 2001; Berggren et al., 2009). The molecular structure of glacial DOM assessed by ultra-high resolution MS has been shown to be more labile than marine or freshwater derived material due to the presence of protein and amino sugar components of microbial origin (Andrilli et al., 2015). This is consistent with studies from other inland waters that show that higher BDOM correlates with enrichment of amino acids and simple sugars (Lafreniere and Sharp, 2004; Williams et al., 2007; Dubnick et al., 2010). A previous comparison of glaciers and rock glaciers (Fegel et al. 2016) used emission excitation matrices (EEMs) to identify an enrichment of proteinaceous DOM in glaciers relative to rock glaciers. In this study, the two compounds that had the greatest difference in abundance between glaciers and rock glaciers, hydroxybenzoic acid (e.g. KEGG C00805, benzenoid superclass) and homoserine (e.g. KEGG C00263, organic acids and derivatives superclass) are present in multiple metabolic pathways including the biosynthesis of siderophore groups and the biosynthesis of secondary metabolites. In addition to the origin of DOM in each glacier type, the pathway the DOM takes to the meltwater outlet also likely differs between glacial types. DOM from rock glaciers may be partially processed by microbes as it travels through the rock glacier’s protosoil. This enhanced “pre-processing” in rock glaciers relative to glaciers may result in DOM with a level of bioavailability more similar to what is delivered to inland surface waters and derived from a terrestrial soil matrix (Fellman et al., 2010). In comparison, glacier BDOM may be preserved in the ice matrix and physically inaccessible to microbial degradation before it is released in glacier meltwater. Thus, the origin of DOM within each glacier type and the pathways through which DOM arrives at outflow are different. These differences in origin and delivery between glacier types are consistent with the differences in the proportion of BDOM between each glacier type that we found in this study. Differences in origin and delivery between glacier types are also consistent with the more similar values between rock glacier BDOM and other surface freshwaters compared to glacier BDOM and other surface waters.

### 4.2 Microbially mediated DOM alteration

Before incubation with a common microbial community, we found that chemical diversity was very similar between glacier and rock glacier DOM (Table 2) and similar to the chemical diversity of DOM found in other freshwater ecosystems (Dubnick et al., 2010; Guillemette and del Giorgio, 2011; Kellerman et al., 2014). However, alteration of DOM during incubation with a common microbial community differed by glacial type. We suggest that differences in a balance between a) the introduction of new metabolites from the common microbial pool relative to b) the loss of BDOM due to microbial consumption may have affected how the diversity of DOM was altered between glacial types during the incubation. For both glacial types incubation with a common microbial community most likely homogenized the DOM pool by eliminating the most labile, or most biologically reactive, compounds and added microbial metabolites through exudates and cellular lysis. As discussed above, rock glacier DOM likely contained more processed DOM (i.e., DOM that had been exposed to diverse microbial communities) and less bioavailable DOM (i.e., secondary metabolites from plants) relative to glacier DOM. Because rock glacier DOM had been more thoroughly exposed to a microbial community before it was collected (relative to glacier DOM), many of the microbial metabolites contributed by the common microbial community were likely already present in rock glacier DOM. Thus, changes in rock glacier DOM composition during the incubation was likely due to loss of labile, biologically reactive components of the DOM pool rather than additions to the DOM pool from the microbial metabolism. This is consistent with the decrease in diversity of the DOM pool we saw for rock glacier DOM. However, the “less processed” glacial DOM may have increased in diversity due to an increase in novel microbial metabolites (relative to the rock glacier DOM) introduced by the common microbial community during the incubation.

The shift in the DOM pool for both glacial types towards compounds provided by microbial metabolism is supported by the increase in the post incubation pool of low molecular weight compounds, often considered to be highly labile. For example, serine, an amino acid, increased between the preincubation and post incubation DOM pools in molecular rank from 1215 to 6, and aspartate from 1049 to 10, and the sugar sucrose increased in molecular rank from 65 to 5 and glycerol from 87 to 4 (Data Set S2), respectively. We expect that microorganisms would rapidly consume these compounds to levels below detection. However, rather than being leftover after the incubation (i.e., recalcitrant compounds inaccessible to microbes), it is more likely that these compounds are an example of the increasing influence of microbial metabolites on the DOM pool during prolonged incubations (i.e., microbial turnover). Compounds in the organic acid superclass which was enriched after incubation include carboxylic acids, amino acids and peptides, which are known products of microbial metabolism. Similarly, purines and fatty acids are part of the organic compound superclass, and carbohydrates are primarily from the organic oxygen superclass. Membership of both classes (organic oxygen and organic compound superclass) also increased over the course of the incubation (Figure 3). This shift in DOM composition towards DOM of microbial origin during the course of the incubation is further supported by an increased similarity of the DOM pool structure (Figure 5) and individual metabolite composition (Figure 6) between the post incubation DOM pool for both glacier types and the alpine lake water control (i.e., experimental control). It is also consistent with the increasing realization that many environmental DOM pools are more likely composed of compounds resulting from microbial metabolism rather than recalcitrant plant metabolites (e.g., Cotrufo et al., 2014). Without a labeled starting material (e.g. an isotopically labeled DOM pool) it is not possible to definitively determine the relative contribution of turnover from microbial metabolism to the post incubation DOM pool. However our results suggest that the post-incubation DOM pool is dominated by products of microbial metabolism.

### 4.3 Analysis of composition and bioavailability of DOM

This study expands the understanding of DOM complexity in inland waters by assessing the glacial DOM inputs to alpine headwaters and moving beyond broad functional groups and compound classes descriptors of DOM to identification of specific compounds, albeit a small (~16%) proportion (Dubnick et al., 2010; Fellman et al., 2010; Singer et al., 2012; Fegel et al., 2016). By using a MS-based technique we were able to identify molecular properties of individual candidate compounds as well as examine a broader swath of chemical diversity from each DOM pool. This approach provides chemical detail at both the individual compound and compositional pool levels. Application of an analytical approach such as mass spectrometry-based metabolomics provides the opportunity to disentangle the critically important components of complex DOM pools for aquatic carbon cycling, however only a few studies have used metabolomic approaches to address the bioavailability of DOM as a heterotrophic subsidy (e.g., see Kujawinski et al., 2011 for a review; Logue et al., 2015).

Although MS based analysis of DOM has important advantages it also comes with a series of challenges and constraints. One challenge comes from the selective DOM extraction using PPL cartridges, which often do not retain very polar compounds (Raeke et al., 2016). However, given the low concentration of DOM (typically ~ 2 mg C per L) in the glacial meltwaters, a concentrating procedure is necessary to perform the GC-MS analyses. Whereas our results provide evidence that differences in bioavailability were concurrent with chemical differences between DOM that differ in origin from glacier types, a significant fraction of the DOM pool was likely not assessed by the GC-MS method we employed. Our GC-MS method scanned for molecules up to 650 *m/z*, leaving larger molecules undetected. Further, many of the components of glacial DOM pools (e.g., the majority of the compounds that differed pre and post incubation) could not be identified using the current databases (NISTv12, Golm, Metlin, and Massbank). Whereas we were unable to determine what those compounds were, we can say what they were not. Primary metabolites are generally well represented in metabolite databases, suggesting that unknown or unannotatable compounds may be products of secondary metabolism rather than primary metabolism, or from input from atmospheric transport of dust and pollutants (Weiland-Brauer et al. 2017). Primary metabolites are considered those metabolically produced compounds that are essential for survival and are also common to many heterotrophic organisms. Secondary metabolites are generally not required for survival, but are organism- or group-specific, and may confer some competitive advantage for an organism that produces them. Secondary metabolites also tend to be more structurally complex than primary metabolites (Luckner, 1984). Products of primary metabolism, e.g. membership in the organic acid, and organic oxygen superclasses are well represented in mass spectrometry spectral libraries and therefore more easily identified with a GC-MS approach. Less well represented are larger molecular weight products of secondary metabolism such with membership in the organoheterocyclic, phenylpropanoid and polyketide superclasses, as well as products of mixed biosynthesis. Known metabolites are often a small portion of data obtained through mass spectrometry (<10%), with much of the data reflecting unknown metabolites or those yet to be verified with standards (Bowen and Northen, 2010). Yet, the quality of mass spectrometry databases and representation is rapidly improving (Blazenovic et al. 2018), and this will likely be a critically important source of information for understanding the relationship between molecular composition of DOM and its bioavailability in future research.

The glaciers and rock glaciers in this study had similar DOM characteristics to other glaciers worldwide and the capability to support heterotrophic metabolism with high (relative to other inland waters) percentages of BDOM in their meltwaters. Our study expands on previous work to characterize the molecular structure of DOM in glacial meltwaters by our novel application of ecosystem metabolomic techniques to glacial meltwater. Whereas we determined that the abundance of 19 compounds differed among glacier types, our results likely only represent a fraction of the chemical compounds controlling the bioavailability of total DOM pools and it is unknown if these compounds would be universally diagnostic for differences in other DOM pools. Other chemical characterization techniques suited towards identifying larger molecules such as liquid chromatography electrospray ionization mass spectrometry (LC-ESI-MS) may provide additional insight into which compounds drive bioavailability because it can access a suite of larger DOM constituents.

## 5. Conclusion

Particularly in the United States, rock glaciers are far more abundant than glaciers in alpine headwater ecosystems (Falaschi et al., 2013; Rangecroft et al., 2015; Fegel et al., 2016). Current glacial carbon modeling neglects the contribution of rock glacial carbon (Hood et al., 2015), yet our results suggest that rock glaciers are a consistent source of BDOM for heterotrophic metabolism in headwater ecosystems. Rock glacier derived DOM may contribute to ecosystem productivity for much longer than glaciers due to the slower recession of rock glaciers compared to glaciers (Woo, 2012; Millar and Westfall, 2010). At similar carbon concentrations and with only ~20% less BDOM (~46% in rock glaciers compared to ~66% in glacier derived DOM), rock glaciers may play just as large if not larger role in heterotrophic metabolism in alpine headwaters as glaciers thaw now and in the near future.

## Acknowledgements

The authors would like to thank Ted Stets and Ann Hess for guidance on the use of the multi-G model. Emily Davidson, and Sarah Lyons helped with interpretation of the metabolomic data. Scott Baggett helped with the statistical analysis, and Gunnar Johnson helped collect DOM samples and create some of the figures used in this manuscript. Gabriel Singer also provided useful feedback on our approach to the statistical analysis of the manuscript. E. K. Hall was supported by NSF award DEB#1456959 during the preparation of this manuscript. A series of anonymous reviewers also improved this manuscript.

## Supporting Information

**Additional Supporting Information (Files uploaded separately)**

Captions for Datasets S1 and S2

## Introduction

This is the supporting information for the manuscript entitled “ASSESSING THE CHEMISTRY AND BIOAVAILABILITY OF DISSOLVED ORGANIC MATTER FROM GLACIERS AND ROCK GLACIERS” by Fegel et al. This document includes two supporting figures, supporting text for methodology, three supporting tables of information, and two .xlsx files of dense samples analysis information. Data was collected from 2014-2015 and was analyzed 2015-2019.

**Text S1:** Derivatization Process for GC-MS

Both pre and post incubation DOM samples were run for gas chromatography-mass spectroscopy (GC-MS) at the Proteomics and Metabolomics Facility at Colorado State University. Pre and post incubation samples were run during the same sample run on the same instrument to ensure instrument relativity. Extracted samples were resuspended in 50 μL of pyridine containing 50 mg mL^−1^ of methoxyamine hydrochloride, incubated at 60°C for 45 min, sonicated for 10 min, and incubated for an additional 45 min at 60°C. 50 μL of N-methyl-N- trimethylsilyltrifluoroacetamide with 1% trimethylchlorosilane (MSTFA + 1% TMCS, Thermo Scientific, Waltham, MA, USA) was added and samples were incubated at 60 °C for 30 min, centrifuged at 3000xg for 5 min, cooled to room temperature, and 80 μL of the supernatant was transferred to a 150 μL glass insert in a GC-MS autosampler vial. Metabolites were detected using a Trace GC Ultra coupled to a Thermo ISQ mass spectrometer (Thermo Scientific, Waltham, MA, USA). Samples were injected in a 1:10 split ratio twice in discrete randomized blocks. Separation occurred using a 30 m TG-5MS column (0.25 mm i.d., 0.25 μm film thickness, Thermo Scientific, Waltham, MA, USA) with a 1.2 mL min^−1^ helium gas flow rate, and the program consisted of 80°C for 30 sec, a ramp of 15°C per min to 330°C, and an 8 min hold. Masses between 50-650 m/z were scanned at 5 scans sec^−1^ after electron impact ionization.

## Data Set S1: Bacterial Data

Please see attached XLSX of all bacterial data used in the study. Starting Cell is the average number of cells per view in the analytical control. Each glacier sample only had bacteria from the analytical control (loch water), therefore we can calculate the number of cells created by comparing the pre to the post. BP is the Bacterial Production rate or the amount of bacterial biomass (as carbon) created per hour. RQ is the respiratory quotient, which is the change in CO2 divided by the change in O2 It is very common practice in limnology to assume an RQ of 1, representative of a heterogenous DOM pool of compounds, not just one specific substrate. Because we did not measure CO2 respiration, this assumption needs to be made. BR is the bacterial respiration rate, or rate that carbon is lost through respiration, using the assumed RQ of 1, multiplied times the rate of dissolved oxygen consumption. BGE or bacterial growth efficiency is the amount of new bacterial biomass produced per unit of organic C substrate assimilated and is a way to relate BP and BR: BGE = (BP)/(BP + BR).

## Data Set S2: GC-MS Data

Metabolite Compounds as Identified through GC-MS, in XLSX format. The “GC-MS Data” tab is the intensities of each compound separated out by individual glacier and rock glacier. The second tab, “Compound Information” is a raw data file of all spectra generated through GC-MS including mass spec contaminates, which were removed prior to the metabolite analysis. Annotation levels (Sumner et al. 2007) are as follows: 1. Identified compounds based on verification with at least two independent and orthogonal data relative to an authentic compound analyzed under identical experimental conditions. 2. Putatively annotated compounds (e.g. without chemical reference standards, based upon physicochemical properties and/or spectral similarity with public/commercial spectral libraries). 3. Putatively characterized compound classes (e.g. based upon characteristic physicochemical properties of a chemical class of compounds, or by spectral similarity to known compounds of a chemical class). 4. Unknown compounds—although unidentified or unclassified these metabolites can still be differentiated and quantified based upon spectral data. Column descriptions are as follows: T

Name: The common chemical name assigned to the compound by the authors.

Original Name: The common chemical name assigned to the compound by the RamSearch software.

Retention Time: The time it takes for chromatogrphic elution from the column in seconds.Confidence Class: A scale of how well the annotated compound fit the library standard compound it was matched to. Annotation confidence are noted with confident 2D structure assignment using a reference standard as a score ‘1’ through a structurally unknown feature of interest of a confidence level ‘4’ (Blazenovic 2018).

Collision Energy: A qualitative descriptor of the intensity of the ion’s collision with the detector. All collision energies were high for our dataset.

Library: The metabolite library used to match the annotated compound.

Library ID: The compound number assigned by the metabolite library.

Match Name:Name of the annotated compound assigned by the metabolite library used to match the compound to a known standard.

InChI Key: International chemical identifier as assigned by the International Union of Pure and Applied Chemistry.

Mass: Molecular mass of the compound, in Daltons.

Match Factor: A numeric value between 1 and 999 identifying how well peaks (both mass and retention time) within the annotated compound matched the peaks within the library compound.

Rev Match Factor: A numeric value between 1 and 999 identifying how well peaks (both mass and retention time) within the annotated compound matched the peaks within the library compound, not including peaks within the sample compound that were not found within the library matched compound. This peak removal helps remove effects of coelution.

Compound Number Peaks: Individual retention times (in seconds) of each of the separated ionized peaks of the compound’s chromatograph.

Intensity Fragment: Intensities of individual ionized peaks within the compound’s chromatograph.

The following are descriptions of each of the ClassyFire superclasses in the dataset transcribed from the Open Biological and Biomedical Ontologies (OBO) format of ChemOnt (http://classyfire.wishartlab.com/downloads): Benzenoid – Aromatic compounds containing one or more benzene rings.

Homogenous non-metal – Inorganic compounds that contain only non-metal elements.

Inorganic – Compounds that do not contain a carbon atom.

Lipids – Fatty acids and their derivatives, and substances related biosynthetically or functionally to these compounds.

Mixed Metal – Compounds that contain both metal and non metal atoms.

Organic Acids – Compounds with an organic acid or derivative thereof.

Organic Carbon – Compounds that contain at least one carbon atom, excluding isocyanide/cyanide and their non-hydrocarbyl derivatives, thiophosgene, carbon diselenide, carbon monosulfide, carbon disulfide, carbon subsulfide, carbon monoxide, carbon trioxide, Carbon suboxide, and dicarbon monoxide.

Organic Nitrogen – Organic compounds containing one or more nitrogen atoms.

Organic Oxygen – Organic compounds containing one or more oxygen atoms.

Organohalogen – Organic compounds containing a bond between a carbon atom and a halogen atom (At, F, Cl, Br, I).

Organoheterocyclic – Compounds containing a ring with least one carbon atom and one non-carbon atom.

Phenylpropanoid – Organic compounds that are synthesized either from the amino acid phenylalanine (phenylpropanoids) or the decarboxylative condensation of malonyl-CoA (polyketides). Phenylpropanoids are aromatic compounds based on the phenylpropane skeleton. Polyketides usually consist of alternating carbonyl and methylene groups (beta-polyketones), biogenetically derived from repeated condensation of acetyl coenzyme A (via malonyl coenzyme A), and usually the compounds derived from them by further condensations.

---

**Figure S1:**
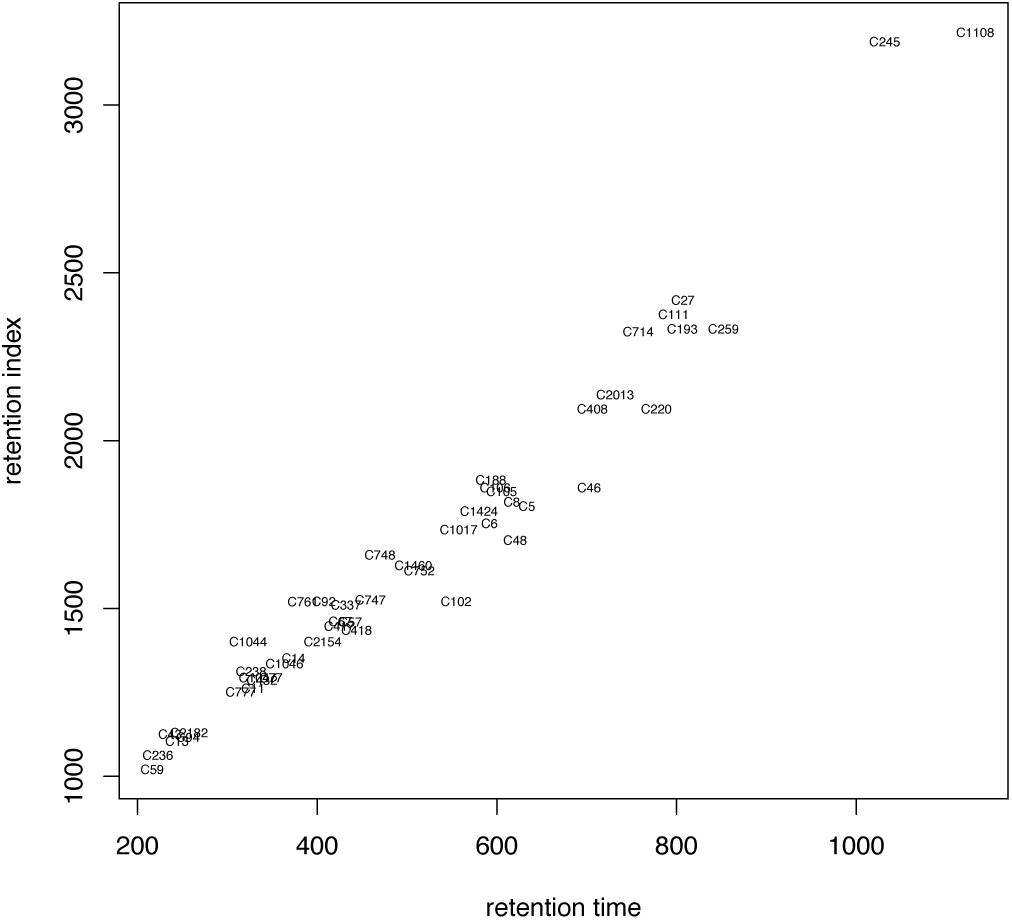
GOLM vs. RT Curve: Comparison of GOLM (metabolite database predicted compound) with those seen with samples on the mass spectrometer at Colorado State University. A linear trend represents an exact match with the annotated compound from the database.

**Figure S2:**
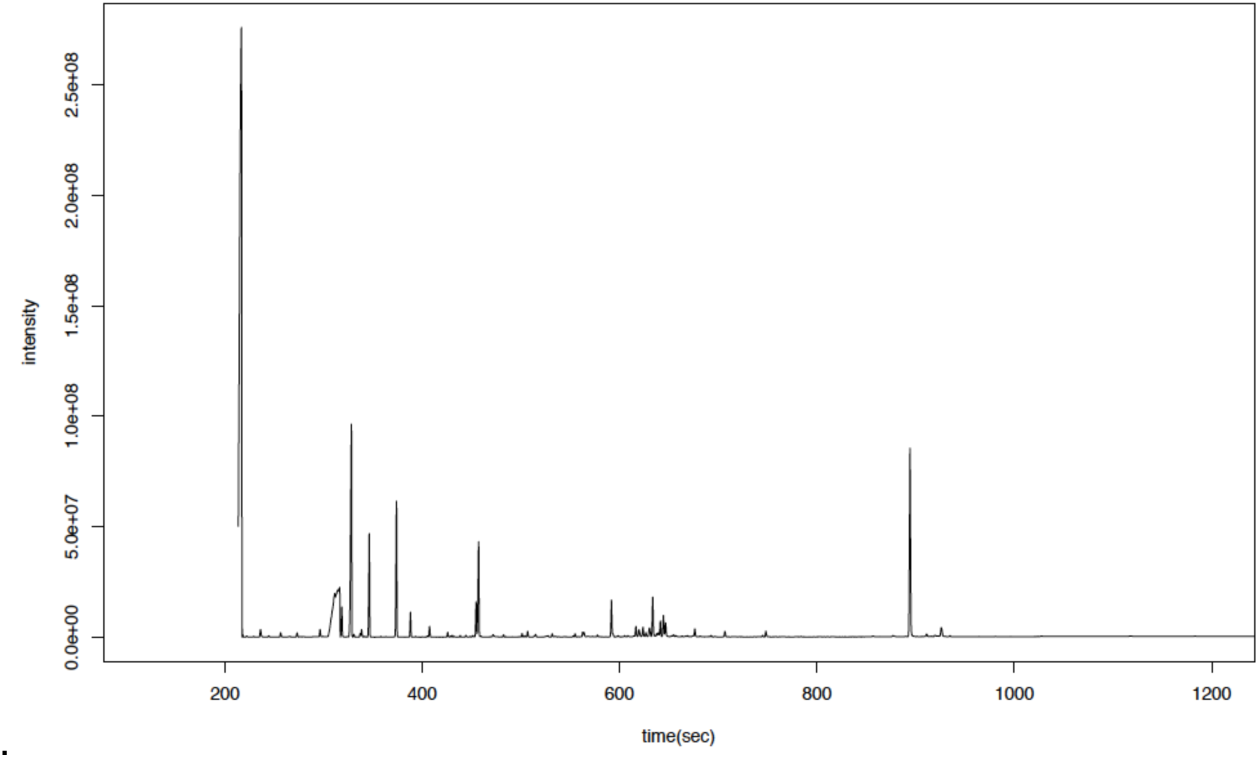
Example chromatogram of GC-MS analyzed compounds in glacial meltwater.

**Table S1:** Standardized BOD Bottle Incubation Recipes.

| SAMPLE NAME | BLANK<br>CORRECTED<br>DOC (MG/L) | TARGET DOC<br>EXPERIMENTAL<br>CONCENTRATION<br>(MG/L) | VOLUME<br>CONCENTRATED<br>DOM ADDED<br>(ML) | VOLUME<br>MILLI-Q<br>ADDED<br>(ML) | TOTAL<br>VOLUME<br>(ML) | VOLUME<br>LOCH<br>WATER<br>ADDED<br>(ML) | MEASURED<br>PRE-INC<br>DOC<br>(MG/L) |
| --- | --- | --- | --- | --- | --- | --- | --- |
| PECK GLACIER | 57.38 | 4.00 | 4.88 | 4.16 | 70 | 60.96 | 4.21 |
| ARAPAHO GLACIER | 40.90 | 4.00 | 6.85 | 2.19 | 70 | 60.96 | 4.09 |
| NAVAJO ROCK<br>GLACIER | 69.29 | 4.00 | 4.04 | 5.00 | 70 | 60.96 | 4.28 |
| ARAPAHO ROCK<br>GLACIER | 73.49 | 4.00 | 3.81 | 5.23 | 70 | 60.96 | 4.42 |
| TAYLOR ROCK<br>GLACIER | 46.03 | 4.00 | 6.08 | 2.95 | 70 | 60.96 | 3.95 |
| ISABELLE GLACIER | 30.99 | 4.00 | 9.04 | 0.00 | 70 | 60.96 | 4.31 |
| ANDREWS GLACIER | 59.19 | 4.00 | 4.73 | 4.31 | 70 | 60.96 | 4.31 |
| PECK ROCK GLACIER | 73.60 | 4.00 | 3.80 | 5.23 | 70 | 60.96 | 4.29 |

**Table S2:** Post Incubation DOC/TDN Data.

| SAMPLE NAME | C CONC. (MG/L) | N CONC.(MG/L) |
| --- | --- | --- |
| ANDREWS GLACIER (REP 1) | 1.28 | 0.53 |
| ANDREWS GLACIER (REP 2) | 1.33 | 0.34 |
| ARAPAHO ROCK GLACIER (REP 1) | 1.03 | 0.46 |
| ARAPAHO GLACIER (REP 1) | 0.89 | 0.42 |
| ARAPAHO GLACIER (REP 2) | 0.79 | 0.52 |
| ARAPAHO GLACIER (REP 2, LAB DUPLICATE) | 0.74 | 0.43 |
| ARAPAHO ROCK GLACIER (REP 2) | 1.09 | 0.43 |
| ISABELLE GLACIER (REP 1) | 0.98 | 0.50 |
| ISABELLE GLACIER (REP 2) | 1.13 | 0.55 |
| EXPERIMENTAL CONTROL (REP 1) | 0.24 | 0.37 |
| EXPERIMENTAL CONTROL (REP 2) | 0.07 | 0.46 |
| FILTER BLANK | 0.02 | 0.07 |
| ANALYTICAL CONTROL | 0.01 | 0.06 |
| NAVAJO ROCK GLACIER (REP 1) | 1.34 | 0.58 |
| NAVAJO ROCK GLACIER (REP2) | 1.17 | 0.56 |
| PECK GLACIER (REP 1) | 1.17 | 0.64 |
| PECK GLACIER (REP 2) | 1.26 | 0.65 |
| PECK ROCK GLACIER (REP 1) | 1.38 | 0.50 |
| PECK ROCK GLACIER (REP 2) | 1.50 | 0.44 |
| TAYLOR ROCK GLACIER (REP 1) | 1.31 | 0.56 |
| TAYLOR ROCK GLACIER (REP 2) | 1.27 | 0.42 |

**Table S3:** DOM Recovery in Pre Incubation Samples.

| SAMPLE NAME | MEASURED<br>CONCENTRATION<br>OF C IN<br>MELTWATER<br>(MG/L) | DOC<br>CONCENTRATION<br>OF EXTRACTED<br>ELUENT (MG/L) | DOM<br>COLLECTED<br>(AS MG OF C ) | CONCENTRATION<br>DOC (MG/L) OF<br>ELUENT PER L OF<br>MELTWATER | DOM<br>RECOVERED<br>(%) |
| --- | --- | --- | --- | --- | --- |
| ANDREWS<br>GLACIER | 0.55 | 16.30 | 1.304 | 0.82 | 66.91 |
| ARAPAHO<br>GLACIER | 0.98 | 21.63 | 1.730 | 1.08 | 90.60 |
| ARAPAHO<br>ROCK GLACIER | 0.34 | 8.40 | 0.672 | 0.42 | 81.94 |
| ISABELLE<br>GLACIER | 0.69 | 15.55 | 1.244 | 0.78 | 89.12 |
| NAVAJO ROCK<br>GLACIER | 0.64 | 24.17 | 1.934 | 1.21 | 52.75 |
| PECK GLACIER | 0.77 | 18.52 | 1.482 | 0.93 | 83.66 |
| PECK ROCK<br>GLACIER | 0.59 | 19.56 | 1.565 | 0.98 | 60.52 |
| TAYLOR ROCK<br>GLACIER | 0.46 | 21.95 | 1.756 | 1.10 | 41.94 |

## References

Amon, Rainer MW, and Ronald Benner. (1996) “Bacterial utilization of different size classes of dissolved organic matter.” Limnology and Oceanography 41.1:41-51.

Anesio, A. M., Hodson, A. J., Fritz, A., Psenner, R., & Sattler, B. (2009). High microbial activity on glaciers: importance to the global carbon cycle. Global Change Biology, 15(4), 955-960.

Antony, R., Sanyal, A., Kapse, N., Dhakephalkar, P. K., Thamban, M., & Nair, S. (2016). Microbial communities associated with Antarctic snow pack and their biogeochemical implications. Microbiological research, 192, 192-202.

Barnes, R. T., Williams, M. W., Parman, J. N., Hill, K., & Caine, N. (2014). Thawing glacial and permafrost features contribute to nitrogen export from Green Lakes Valley, Colorado Front Range, USA. Biogeochemistry, 117(2-3), 413-430.

Baron, J.S., Fegel, T., Boot, C.M., and Hall, E.K., 2019, Laboratory Incubation results from 2015 for bacterial cell counts, carbon use efficiency, growth efficiency, and dissolved organic matter chemistry from four glacier outflows and four rock glacier outflows in Colorado: U.S. Geological Survey data release, https://doi.org/10.5066/P9A0JNP9.

Benner, Ronald. (2002) “Chemical composition and reactivity.” Biogeochemistry of marine dissolved organic matter 3: 56-90.

Berggren, Martin, and Paul A. Giorgio. (2015) “Distinct patterns of microbial metabolism associated to riverine dissolved organic carbon of different source and quality.” Journal of Geophysical Research: Biogeosciences.

Berner, Robert A. (1980) Early diagenesis: A theoretical approach. No. 1. Princeton University Press.

Berthling, Ivar. (2011) “Beyond confusion: Rock glaciers as cryo-conditioned landforms.” Geomorphology 131.3: 98-106.

Blaženović, I., Kind, T., Ji, J., & Fiehn, O. (2018). Software tools and approaches for compound identification of LC-MS/MS data in metabolomics. Metabolites, 8(2), 31.

Borsheim, Knut Yngve, and Gunnar Bratbak. (1987) “Cell volume to cell carbon conversion factors for a bacterivorous Monas sp. enriched from seawater.” Mar. Ecol. Prog. Ser 36.17: ll.

Bowen, Benjamin P., and Trent R. Northen. (2010) “Dealing with the unknown: metabolomics and metabolite atlases.” Journal of the American Society for Mass Spectrometry 21.9: 1471-1476.

Broeckling, C. D., Afsar, F. A., Neumann, S., Ben-Hur, A., & Prenni, J. E. (2014). RAMClust: a novel feature clustering method enables spectral-matching-based annotation for metabolomics data. Analytical chemistry, 86(14), 6812-6817.

Burga, C. A., Frauenfelder, R., Ruffet, J., Hoelzle, M., & KääB, A. N. D. R. E. A. S. (2004). Vegetation on Alpine rock glacier surfaces: a contribution to abundance and dynamics on extreme plant habitats. Flora-Morphology, Distribution, Functional Ecology of Plants, 199(6), 505-515.

Cannone, Nicoletta, and Renato Gerdol. (2003) “Vegetation as an ecological indicator of surface instability in rock glaciers.” Arctic, Antarctic, and Alpine Research 35.3: 384-390.

Cotrufo, MF, Wallenstein, MD, Boot, CM, Denef, K Paul, E. (2014) “The Microbial Efficiency Matrix Stabilization (MEMS) framework integrates plant litter decomposition with soil organic matter stabilization: do labile plant inputs form stable soil organic matter?” Global Change Biology 19 (4): 988-995

D’Andrilli, J., Cooper, W. T., Foreman, C. M., & Marshall, A. G. (2015). An ultrahigh-resolution mass spectrometry index to estimate natural organic matter lability. Rapid Communications in Mass Spectrometry, 29(24), 2385-2401.

Del Giorgio, Paul A., and Jonathan J. Cole. (1998) “Bacterial growth efficiency in natural aquatic systems.” Annual Review of Ecology and Systematics: 503-541.

Derenne, Sylvie, and Thanh Thuy Nguyen Tu. (2014) “Characterizing the molecular structure of organic matter from natural environments: An analytical challenge.” Comptes Rendus Geoscience 346.3: 53-63.

Dittmar, T., Koch, B., Hertkorn, N., & Kattner, G. (2008). A simple and efficient method for the solid-phase extraction of dissolved organic matter (SPE-DOM) from seawater. Limnology and Oceanography: Methods, 6(6), 230-235.

Dubnick, A., Barker, J., Sharp, M., Wadham, J., Lis, G., Telling, J., … & Jackson, M. (2010). Characterization of dissolved organic matter (DOM) from glacial environments using total fluorescence spectroscopy and parallel factor analysis. Annals of Glaciology, 51(56), 111-122.

Falaschi, D., Bravo, C., Masiokas, M., Villalba, R., & Rivera, A. (2013). First glacier inventory and recent changes in glacier area in the Monte San Lorenzo Region (47 S), Southern Patagonian Andes, South America. Arctic, Antarctic, and Alpine Research, 45(1), 19-28.

Fegel, T. S., Baron, J. S., Fountain, A. G., Johnson, G. F., & Hall, E. K. (2016). The differing biogeochemical and microbial signatures of glaciers and rock glaciers. Journal of Geophysical Research: Biogeosciences, 121(3), 919-932.

Fellman, Jason B., Eran Hood, and Robert GM Spencer. (2010) “Fluorescence spectroscopy opens new windows into dissolved organic matter dynamics in freshwater ecosystems: A review.” Limnology and Oceanography 55.6: 2452-2462.

Fellman, J. B., Spencer, R. G., Hernes, P. J., Edwards, R. T., D’Amore, D. V., & Hood, E. (2010). The impact of glacier runoff on the biodegradability and biochemical composition of terrigenous dissolved organic matter in near-shore marine ecosystems. Marine Chemistry, 121(1-4), 112-122.

Fellman, J. B., Hood, E., Raymond, P. A., Hudson, J., Bozeman, M., & Arimitsu, M. (2015). Evidence for the assimilation of ancient glacier organic carbon in a proglacial stream food web. Limnology and Oceanography, 60(4), 1118-1128.

Feunang, Y. D., Eisner, R., Knox, C., Chepelev, L., Hastings, J., Owen, G., … & Greiner, R. (2016). ClassyFire: automated chemical classification with a comprehensive, computable taxonomy. Journal of cheminformatics, 8(1), 61.

Guenet, B., Danger, M., Abbadie, L., & Lacroix, G. (2010). Priming effect: bridging the gap between terrestrial and aquatic ecology. Ecology, 91(10), 2850-2861.

Guillemette, François, and Paul A. del Giorgio. (2011) “Reconstructing the various facets of dissolved organic carbon bioavailability in freshwater ecosystems. “Limnology and Oceanography 56.2: 734-748.

Halket, J. M., Waterman, D., Przyborowska, A. M., Patel, R. K., Fraser, P. D., & Bramley, P. M. (2004). Chemical derivatization and mass spectral libraries in metabolic profiling by GC/MS and LC/MS/MS. Journal of experimental botany, 56(410), 219-243.

Hobbie, J. El. R. Jasper Daley, and STTI977 Jasper. (1977) “Use of nucleopore filters for counting bacteria by fluorescence microscopy.” Applied and environmental microbiology 33.5: 1225-1228.

Hood, E., Fellman, J., Spencer, R. G., Hernes, P. J., Edwards, R., D’Amore, D., & Scott, D. (2009). Glaciers as a source of ancient and labile organic matter to the marine environment. Nature, 462(7276), 1044.

Hood, E., Battin, T. J., Fellman, J., O’neel, S., & Spencer, R. G. (2015). Storage and release of organic carbon from glaciers and ice sheets. Nature geoscience, 8(2), 91.

Keller, B. O., Sui, J., Young, A. B., & Whittal, R. M. (2008). Interferences and contaminants encountered in modern mass spectrometry. Analytica chimica acta, 627(1), 71-81.

Kellerman, Anne M., et al. (2014) “Chemodiversity of dissolved organic matter in lakes driven by climate and hydrology.” Nature communications 5.

Kellerman, A. M., Dittmar, T., Kothawala, D. N., & Tranvik, L. J. (2014). Chemodiversity of dissolved organic matter in lakes driven by climate and hydrology. Nature communications, 5, 3804.

Kind, Tobias, and Oliver Fiehn. (2010) “Advances in structure elucidation of small molecules using mass spectrometry.” Bioanalytical reviews 2.1-4: 23-60.

Kujawinski, E. B., Freitas, M. A., Zang, X., Hatcher, P. G., Green-Church, K. B., & Jones, R. B. (2002). The application of electrospray ionization mass spectrometry (ESI MS) to the structural characterization of natural organic matter. Organic geochemistry, 33(3), 171-180.

Kujawinski, Elizabeth B. (2011) “The impact of microbial metabolism on marine dissolved organic matter.” Annual review of marine science 3: 567-599.

Lafrenière, Melissa J., and Martin J. Sharp. (2004) “The concentration and fluorescence of dissolved organic carbon (DOC) in glacial and nonglacial catchments: interpreting hydrological flow routing and DOC sources.” Arctic, Antarctic, and Alpine Research 36.2: 156-165.

Logue, J. B., Stedmon, C. A., Kellerman, A. M., Nielsen, N. J., Andersson, A. F., Laudon, H., … & Kritzberg, E. S. (2016). Experimental insights into the importance of aquatic bacterial community composition to the degradation of dissolved organic matter. The ISME journal, 10(3), 533.

Luckner, Martin. (1984) “Secondary Metabolism, a Distinct Part of General Metabolism.” Secondary Metabolism in Microorganisms, Plants and Animals. Springer Berlin Heidelberg. 25-30.

Millar, Constance I., and Robert D. Westfall. (2010) “Distribution and climatic relationships of the American pika (Ochotona princeps) in the Sierra Nevada and western Great Basin, USA; periglacial landforms as refugia in warming climates.” Arctic, Antarctic, and Alpine Research 4

Mostovaya, A., Hawkes, J. A., Koehler, B., Dittmar, T., & Tranvik, L. J. (2017). Emergence of the reactivity continuum of organic matter from kinetics of a multitude of individual molecular constituents. Environmental science & technology, 51(20), 11571-11579.

Presens Oxy-4 Mini: 4-Channel Fiber optic oxygen transmitter Instruction Manual (2011) Precision Sensing GmbH p. 28-31

Pierce, Alan Eugene. (1968) “Silylation of organic compounds.”

Raeke, J., Lechtenfeld, O. J., Wagner, M., Herzsprung, P., & Reemtsma, T. (2016). Selectivity of solid phase extraction of freshwater dissolved organic matter and its effect on ultrahigh resolution mass spectra. Environmental Science: Processes & Impacts, 18(7), 918-927.

Raes, Jeroen and Peer Bork. (2008) “Molecular eco-systems biology: toward and understanding of community function.” Nature Reviews Microbiology 6: 693-699.

Raymond, Peter A., and James E. Bauer. (2001) “Riverine export of aged terrestrial organic matter to the North Atlantic Ocean.” Nature 409.6819: 497.

R Core Team (2014). R: A language and environment for statistical computing. R Foundation for Statistical Computing, Vienna, Austria. URL http://www.R-project.org/.

Rangecroft, S., S. Harrison, and K. Anderson. (2015) “Rock glaciers as water stores in the Bolivian Andes: an assessment of their hydrological importance.” Arctic, Antarctic, and Alpine Research 47.1: 89-98.

Rovira, Pere, and V. Ramón Vallejo. (2002) “Labile and recalcitrant pools of carbon and nitrogen in organic matter decomposing at different depths in soil: an acid hydrolysis approach.” Geoderma 107.1: 109-141.

Sanyal, A., Antony, R., Samui, G., & Thamban, M. (2018). Microbial communities and their potential for degradation of dissolved organic carbon in cryoconite hole environments of Himalaya and Antarctica. Microbiological research, 208, 32-42.

Saros, J. E., Rose, K. C., Clow, D. W., Stephens, V. C., Nurse, A. B., Arnett, H. A., … & Wolfe, A. P. (2010). Melting alpine glaciers enrich high-elevation lakes with reactive nitrogen. Environmental science & technology, 44(13), 4891-4896.

Shannon, C. E. (1948) “A Mathematical Theory of Communication-An Integrated Approach.”.

Singer, G. A., Fasching, C., Wilhelm, L., Niggemann, J., Steier, P., Dittmar, T., & Battin, T. J. (2012). Biogeochemically diverse organic matter in Alpine glaciers and its downstream fate. Nature Geoscience, 5(10), 710.

Slemmons, Krista EH, Jasmine E. Saros, and Kevin Simon. (2013) “The influence of glacial meltwater on alpine aquatic ecosystems: a review.” Environmental Science: Processes & Impacts 15.10: 1794-1806.

Spencer, R. G., Guo, W., Raymond, P. A., Dittmar, T., Hood, E., Fellman, J., & Stubbins, A. (2014). Source and biolability of ancient dissolved organic matter in glacier and lake ecosystems on the Tibetan Plateau. Geochimica et Cosmochimica Acta, 142, 64-74.

Stibal, M., Lawson, E. C., Lis, G. P., Mak, K. M., Wadham, J. L., & Anesio, A. M. (2010). Organic matter content and quality in supraglacial debris across the ablation zone of the Greenland ice sheet. Annals of Glaciology, 51(56), 1-8.

Stubbins, A., Hood, E., Raymond, P. A., Aiken, G. R., Sleighter, R. L., Hernes, P. J., … & Abdulla, H. A. (2012). Anthropogenic aerosols as a source of ancient dissolved organic matter in glaciers. Nature Geoscience, 5(3), 198.

Sumner, L. W., Amberg, A., Barrett, D., Beale, M. H., Beger, R., Daykin, C. A., … & Hankemeier, T. (2007). Proposed minimum reporting standards for chemical analysis. Metabolomics, 3(3), 211-221.

Sun, L., Perdue, E. M., Meyer, J. L., & Weis, J. (1997). Use of elemental composition to predict bioavailability of dissolved organic matter in a Georgia river. Limnology and Oceanography, 42(4), 714-721.

Vachon, D., Prairie, Y. T., Guillemette, F., & Del Giorgio, P. A. (2017). Modeling allochthonous dissolved organic carbon mineralization under variable hydrologic regimes in boreal lakes. Ecosystems, 20(4), 781-795.

Vähätalo, Anssi V., Hanna Aarnos, and Samu Mäntyniemi. “Biodegradability continuum and biodegradation kinetics of natural organic matter described by the beta distribution.” Biogeochemistry 100.1-3 (2010): 227-240.

Volk, Christian J., Catherine B. Volk, and Louis A. Kaplan. (1997) “Chemical composition of biodegradable dissolved organic matter in streamwater.” Limnology and Oceanography 42.1: 39-44.

Wahrhaftig, Clyde, and Allan Cox. (1959) “Rock glaciers in the Alaska Range.” Geological Society of America Bulletin 70.4: 383-436.

Wetzel, Robert G. (1992) “Gradient-dominated ecosystems: sources and regulatory functions of dissolved organic matter in freshwater ecosystems.” Dissolved organic matter in lacustrine ecosystems. Springer Netherlands. 181-198.

Wiegner, T. N., Seitzinger, S. P., Glibert, P. M., & Bronk, D. A. (2006). Bioavailability of dissolved organic nitrogen and carbon from nine rivers in the eastern United States. Aquatic Microbial Ecology, 43(3), 277-287.

Weiland-Bräuer, N., Fischer, M. A., Schramm, K. W., & Schmitz, R. A. (2017). Polychlorinated biphenyl (PCB)-degrading potential of microbes present in a cryoconite of Jamtalferner glacier. Frontiers in microbiology, 8, 1105.

Williams, M. W., Knauf, M., Caine, N., Liu, F., & Verplanck, P. L. (2006). Geochemistry and source waters of rock glacier outflow, Colorado Front Range. Permafrost and Periglacial Processes, 17(1), 13-33.

Williams, M. W., Knauf, M., Cory, R., Caine, N., & Liu, F. (2007). Nitrate content and potential microbial signature of rock glacier outflow, Colorado Front Range. Earth Surface Processes and Landforms: The Journal of the British Geomorphological Research Group, 32(7), 1032-1047.

Woo, Ming-ko. (2012) Permafrost hydrology. Springer Science & Business Media.

